# CroCoNet: a novel framework for cross-species network analysis reveals POU5F1 (OCT4) rewiring

**DOI:** 10.1101/2025.11.18.689002

**Authors:** Anita Térmeg, Vladyslav Storozhuk, Zane Kliesmete, Fiona C. Edenhofer, Johanna Geuder, Beate Vieth, Philipp Janssen, Ines Hellmann

**Author notes:** correspondence: Ines Hellmann, Telefon, www.anthropologie.bio.lmu.de.

## Abstract

**BACKGROUND:** Gene regulatory changes play a central role in shaping cellular phenotypes across species. To understand how these phenotypes evolve, it is essential to investigate the underlying gene regulatory networks (GRNs). However, most comparative analyses of GRNs remain qualitative and are therefore sensitive to false positives and false negatives.

**RESULTS:** To address this limitation, we introduce CroCoNet (**Cro**ss-species **Co**mparison of **Net**works), an R package for the quantitative comparison of GRNs across species. CroCoNet constructs comparable network modules centered on known transcriptional regulators and quantifies the variability in module topology within and between species. By contrasting these levels of variability, CroCoNet can distinguish true evolutionary divergence from technical and biological confounders. Applying CroCoNet to scRNA-seq data from the early neural differentiation of human, gorilla, and cynomolgus macaque, we identified 20 conserved and 24 diverged modules. Despite the conserved expression pattern of the pluripotency factor POU5F1 (OCT4), its associated module was among the most diverged. This result was independently confirmed through cross-species CRISPRi perturbations coupled with single-cell RNA-seq as a readout. Moreover, we found that great ape- and human-specific LTR7 elements are enriched near POU5F1 module genes, potentially contributing to the cross-species differences in network topology.

**CONCLUSIONS:** These findings demonstrate that CroCoNet can resolve regulatory rewiring and provides a robust framework for studying GRN evolution across closely related species. CroCoNet is available as an open-source R package at https://hellmann-lab.github.io/CroCoNet/.

## Background

Linking genotype to phenotype is a central goal in evolutionary biology. Gene regulatory networks (GRNs), which coordinate transcriptional programs across the genome, are key intermediaries in this relationship. Studying GRNs at the level of co-expression modules provides a powerful abstraction: modules often reflect shared regulatory inputs or common functional roles, and can be more conserved across species than individual genes [1, 2]. Module-based comparative analyses of GRNs therefore offer insights into the molecular basis of phenotypic evolution, while at the same time informing more accurate models of regulatory network organization.

Understanding network evolution is an essential step in connecting our knowledge of molecular evolution to the observed phenotypic diversity, yet it remains relatively poorly characterized in vertebrates. While microorganisms have been studied extensively, large-genome organisms with smaller effective population sizes may follow different evolutionary rules. In mammals, most cross-species comparisons have focused on evolutionarily distant species such as humans and mice [3], where regulatory differences are dominated by gene dosage changes. However, given the architecture of mammalian genomes, changes in transcription factor activity and modifications to cis-regulatory elements are likely to represent more immediate and widespread sources of regulatory evolution than dosage alone. To study these processes, it is advantageous to focus on module divergence among closely related species.

When discussing module conservation, it is important to distinguish between two conceptually different aspects. One is the conservation of a module’s overall expression pattern (often summarized by its “eigengene”), which reflects the combined activity of the genes in the module [4–6]. The other aspect is the conservation of the module’s network topology, defined by the pattern and strength of regulatory relationships among its member genes [7, 8]. These two aspects can diverge: a module may retain a similar expression profile across species even while its internal regulatory wiring changes. Such topological changes can be compensatory, maintaining similar functional outputs despite underlying mechanistic rewiring, a phenomenon called developmental system drift [9]. In the present work, we focus on the conservation and divergence of module topology.

The emergence of single-cell RNA sequencing (scRNA-seq) technologies has made it possible to study GRNs at an unprecedented scale and resolution. Profiling thousands to millions of individual cells in a single experiment provides sufficient observations for the detection of co-expression patterns [10]. Moreover, analyzing single cells instead of bulk cell type mixtures is bound to provide more meaningful networks [11, 12]. However, measures of expression similarity are still affected by substantial biological and technical noise, including dropouts, background contamination, and variation in sequencing depth [13–15].

This low signal-to-noise ratio is especially challenging in a cross-species setting. A common strategy for comparing GRNs across species is to define modules independently in each species and quantify their overlap. However, this approach is highly sensitive to false positives and false negatives in module detection. Even if the true underlying modules are identical, limited power (e.g., ∼50% in each species) already reduces the observed overlap to ∼33%. False positives further reduce the perceived conservation, as they rarely overlap, especially for small modules. Because error rates also depend on gene expression levels [13–15], highly expressed regulators may appear more conserved. A fair evolutionary comparison, therefore, requires (1) more robust statistical measures than membership overlap, and (2) an explicit adjustment for the sources of variance that underlie differences in power.

Our software, **Cro**ss-species **Co**mparison of **Net**works (CroCoNet), is designed to meet both of these requirements. To address the first, we incorporate module preservation statistics to compare module topologies across species. To address the second, we use variation observed across biological replicates to correct the preservation scores for confounding factors.

## Results

### Overview of the CroCoNet pipeline

CroCoNet consists of three main steps: 1) identifying consensus modules by integrating information from all species, 2) comparing module topology across species as well as across replicates of the same species, and 3) quantifying module conservation by contrasting the degree of topological differences within versus across species (Fig. 1).

**Figure 1.**
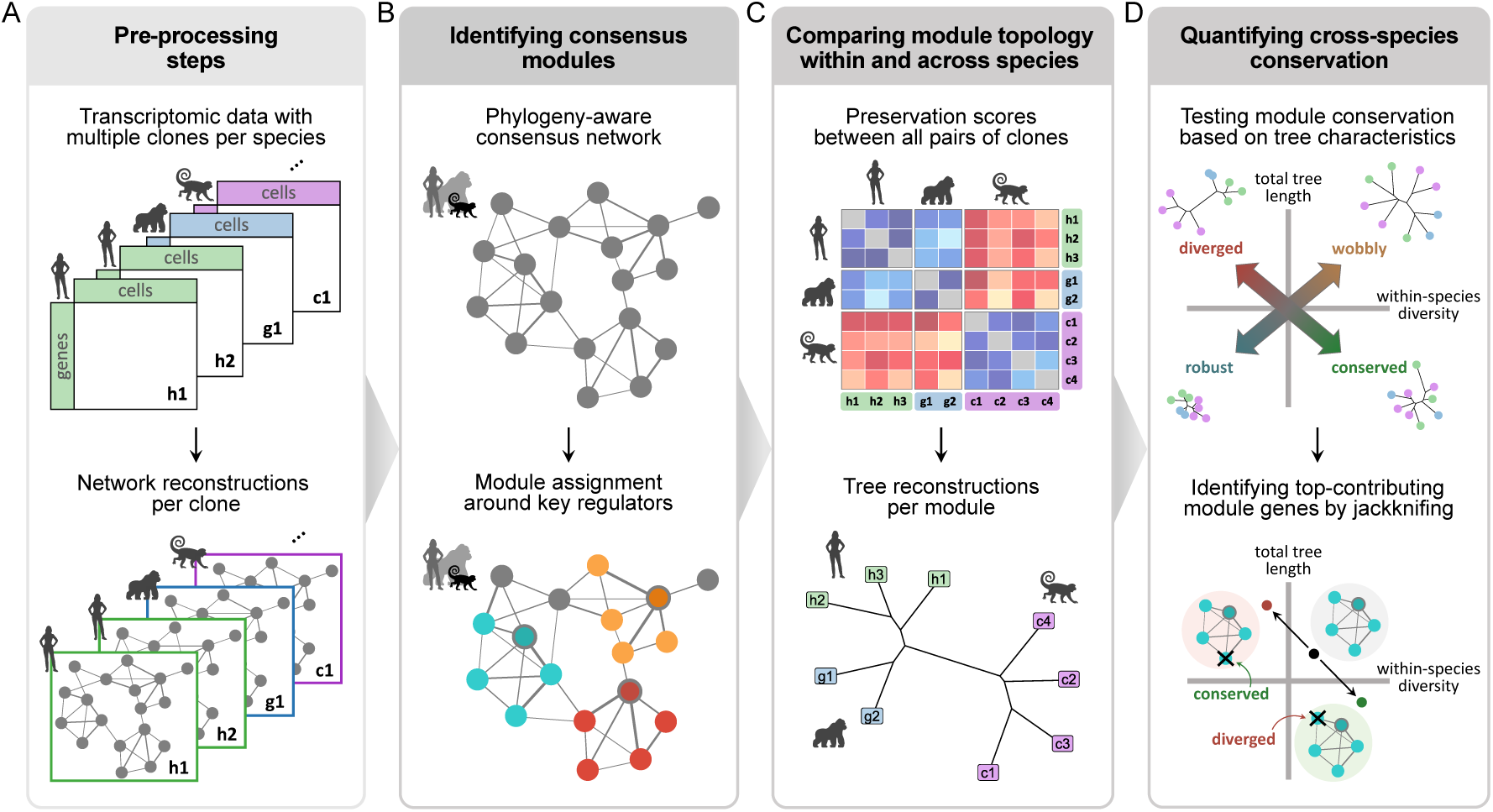
Overview of the CroCoNet pipeline. **A**) The input for the pipeline is one network reconstruction per biological replicate. **B**) The networks are combined into a single consensus network across replicates and species. Co-expression modules of chosen key regulators are assigned based on the consensus network. **C**) The modules are compared between all replicates using a preservation score [7]. The preservation scores are turned into a distance matrix that is used to summarize module preservation as a neighbor-joining tree. **D**) Contrasting the total tree length with the sum of all within-species branch lengths provides a power-corrected conservation measure. Jackknifing the modules provides a metric for robustness and also allows us to identify influential target genes that contribute the most to the module conservation or divergence.

CroCoNet is designed for the analysis of scRNA-seq data from multiple species with multiple biological replicates per species. The experimental conditions, pseudotime trajectory and cell type composition should be as comparable across species as possible to ensure meaningful results. In addition, the analyzed gene set needs to be matched across species to retain only one-to-one correspondences. These well-matched data are the basis for network analysis.

The input for CroCoNet is a set of networks, one for each biological replicate (Fig. 1A). Network inference must therefore be performed prior to running CroCoNet, using any method of choice that produces adjacency measures. In our examples, we used two methods: GRNBoost2 [16], a gradient boosting-based approach that performed well in multiple benchmarks [14, 17], and Spearman’s correlation, a simple and computationally efficient alternative. Users must also provide a list of putative regulators, which can be defined based on prior knowledge, data-driven criteria, or a combination of the two. In our examples, we used genes annotated as transcriptional regulators (TRs) [18, 19] that also showed above-noise expression variance in at least one species of our dataset.

The CroCoNet pipeline begins with the identification of consensus co-expression modules (Fig. 1B). To this end, replicate-wise networks are first combined into a consensus network. Consensus modules are then assigned around the provided regulators based on regulator–target adjacencies (i.e., how strongly a gene is connected to the central regulator) and intramodular connectivities (i.e., how strongly a gene is connected to the rest of the module). Our module assignment uses a dynamic filtering procedure, in which modules must stay above a minimum gene count but are otherwise free to vary in size.

The second step is to compare each module between all pairs of replicates, both within and across species (Fig. 1C). To this end, we adapted two preservation statistics suggested by Langfelder *et al.*: the correlation of adjacencies and the correlation of intramodular connectivities [7]. Both are designed to quantify the degree of topological similarity between two networks but strike a different balance between sensitivity and specificity. These pairwise preservation statistics are converted into a distance matrix which is then summarized as a neighbor-joining tree.

In the final step, tree-based statistics are used to derive a metric of evolutionary conservation/divergence (Fig. 1D). In the spirit of many tests commonly used to infer selection from sequence variation [20, 21], we contrast differences within and between species to estimate the rate of meaningful changes. In the context of network analysis, within-species diversity is shaped not only by genetic differences across individuals but also to a large extent by environmental and technical noise. The sum total of this — what we measure here as within-species diversity — can be so large that it might obscure true between-species differences. Fitting a linear model with the total tree length as dependent and within-species diversity as independent variable yields a slope that informs us about the average contribution of between-species differences to total network variability. The residuals of the linear regression model reflect deviations from this average contribution and can thus be used as measures of module divergence.

### Applying CroCoNet to primate scRNA-seq data

To demonstrate the utility of CroCoNet, we applied it to two primate cross-species scRNA-seq datasets. The first dataset, which was generated in our lab, profiles early neural differentiation in human, gorilla, and cynomolgus macaque. In addition, we analyzed a published dataset comparing brain samples from five primate species [22, 23] (Supplementary Fig. S1). Throughout the manuscript, we focus on the neural differentiation data as the main example, and provide complementary insights from the brain data.

To obtain the main example dataset, three iPSC clones from three humans, two iPSC clones from one gorilla, and four iPSC clones from two cynomolgus macaque individuals (Supplementary Table 1) were differentiated into neural progenitor cells (NPCs). During the 9-day differentiation process, cells were profiled using scRNA-seq at six different time points (Fig. 2A). To create a shared gene space for counting, we transferred the human GENCODE gene models [24] to the gorilla and cynomolgus macaque genomes using Liftoff [25]. Note that this approach is only feasible for closely related species and only makes sense when the reference gene models are superior [26]. After the preprocessing steps, pseudotime analysis of the ∼4,000 cells revealed a clear trajectory from pluripotent stem cells to early ectoderm and neurons (Fig. 2B) that aligned well across the three species (Supplementary Fig. S2A). With comparability across species confirmed, we used GRNBoost2 [16] to reconstruct fully connected GRNs per replicate (Supplementary Fig. S2B, see Methods).

**Figure 2.**
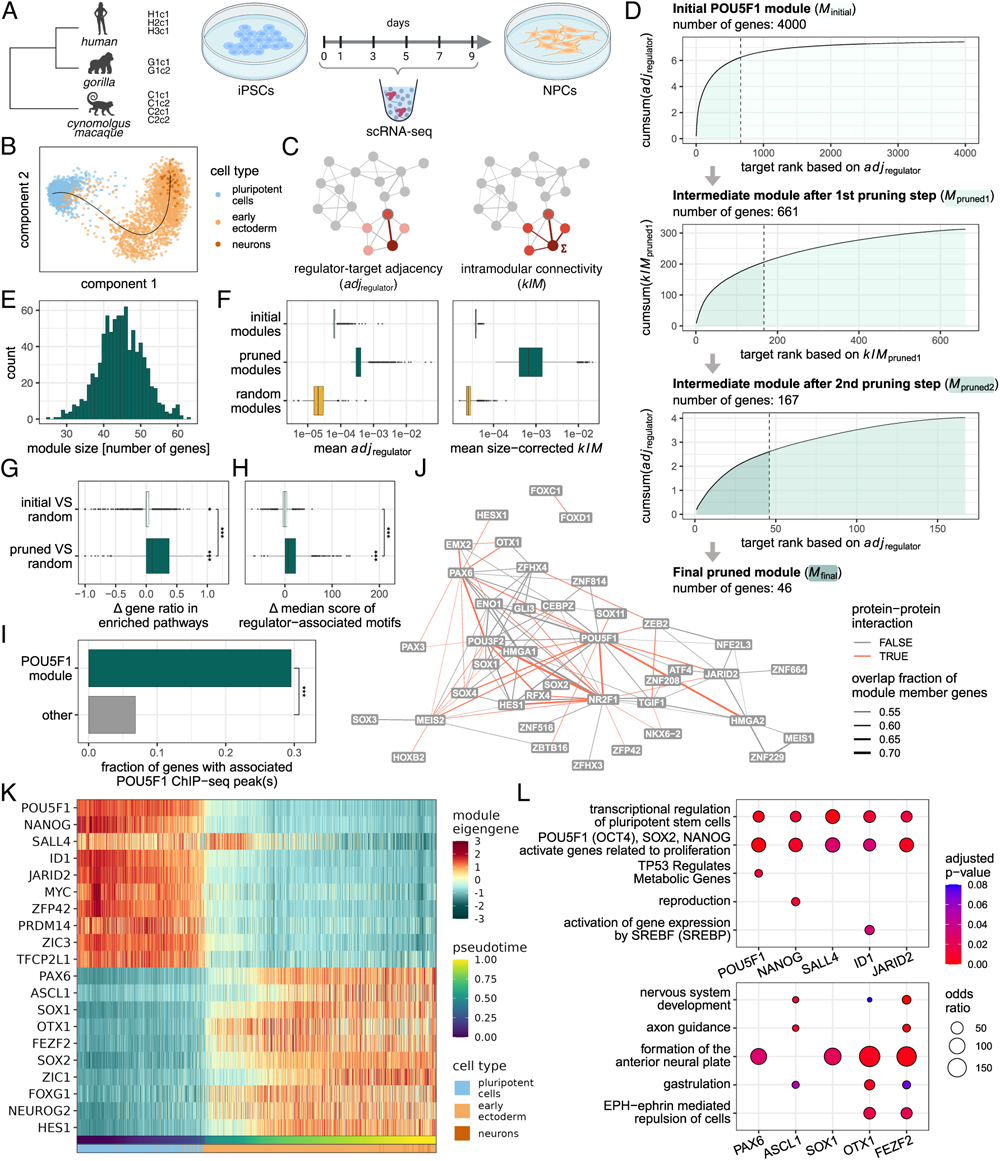
Assigning biologically meaningful consensus modules. **A**) Experimental setup of the scRNA-seq experiment. **B**) Pseudotime trajectory with cells colored by cell type. **C**) Calculation of the regulator–target adjacency (*adj_regulator_*) and the intramodular connectivity (kIM), the statistics used to prune the modules. **D**) Dynamic module pruning, exemplified by the POU5F1 module. We iteratively calculate the cumulative sum curve of *adj_regulator_*or *kIM* and keep targets that ranked higher than the knee point of the curve. **E**) Distribution of module sizes after pruning. **F**) Mean *adj*_regulator_ and mean size-corrected *kIM* (module-level summaries of the statistics underlying pruning) of the initial, pruned and random modules. **G**) Pathway enrichment was quantified per module as the fraction of genes associated with any enriched Reactome pathway. The plot shows how the enrichment in the initial and pruned modules differs relative to the random modules. **H**) Each target gene was scored for binding motifs of the module’s central regulator in the human genome (see Methods). The plot shows how the module-level summaries of these motif scores differ in the initial and pruned modules relative to the random modules. **I**) POU5F1 ChIP-seq peaks [27] are enriched near POU5F1 module genes. **J**) Module overlaps and protein–protein interactions of the regulators. Nodes represent modules associated with the regulators on the label and edge weights represent the overlap fraction of module members. Only the 100 highest overlaps are shown. Red edges indicate that the regulators interact based on STRINGdb [28]. **K**) Pseudotime trajectories of the module eigengenes for 10 pluripotency [29–34] and 10 early neural markers [35–40]. The eigengene calculation was based on the positively correlated targets. **L**) Pathways with the highest enrichment across the modules of 5 pluripotency markers (top) and 5 neural differentiation markers (bottom). **G–I**) Asterisks indicate significance: * *p <* 0.05, ** *p <* 0.01, *** *p <* 0.001. _9_

For the brain data, we used the published count matrices and cell type annotations distinguishing 18 neural and 6 glial subclasses. Prior to network inference, we subsampled all replicates to the same number of cells with the same cell type proportions, resulting in ∼9,500 cells per replicate (Supplementary Table 2). In the case of this larger dataset, we used gene-gene Spearman’s correlations to infer replicate-wise networks.

In both cases, these preprocessing steps produced one undirected, unsigned GRN per biological replicate, and these sets of networks served as input for the CroCoNet pipeline.

### Consensus network and module assignment

The first step of CroCoNet is to assign consensus modules jointly for all species. The goal is to create modules that are unbiased and meaningful – unbiased in the sense that edges are equally likely to be detected in all species and meaningful in that they are enriched for the actual targets of the regulator.

To achieve this, replicate-wise GRNs were combined into one consensus GRN. To avoid biasing the consensus network, adjacencies from all replicates were weighted according to the species’ phylogeny and the number of replicates before taking the average. Based on these consensus adjacencies, more specifically based on the adjacency of a gene to one of the 836 putative regulators (Supplementary Fig. S2C-D), we assigned initial modules, each containing 4,000 genes. These modules were then pruned using a dynamic pruning approach that iteratively selects genes based on two metrics: the regulator–target adjacency and the intramodular connectivity (Fig. 2C). In each pruning step, we calculated the cumulative sum curves based on one of these two metrics per module, then kept the targets ranking higher than the knee point of the curve (Fig. 2D). We continued the iterations as long as all modules contained at least 20 genes, resulting in a median module size of 45 (Fig. 2E, Supplementary Tables 3–4).

Note that, in contrast to WGCNA [41], from which we adapted the preservation statistics, module memberships in CroCoNet are not mutually exclusive. We consider this important, as a realistic representation of gene regulation should also capture the cooperative nature of TRs. Using protein–protein interaction data from STRINGdb [28], we confirmed that interacting regulators in our network tend to share more module member genes than non-interacting ones (Wilcoxon test, *n*_1_ = 17, 670, *n*_2_ = 331, 360, *p <* 1 · 10^−16^, Fig. 2J, Supplementary Fig. S3A-C). This highlights that CroCoNet effectively captures cooperative regulatory mechanisms.

To assess the biological relevance of the consensus modules, we first examined the enrichment of module genes in known biological pathways. As a reference point for all such module evaluations, we created random modules alongside the actual modules. To this end, we drew target genes at random to match the size of the corresponding pruned module for each regulator (Fig. 2F). The pruned modules contained a significantly higher proportion of genes associated with enriched Reactome pathways [42, 43] compared to the random or initial modules (paired Wilcoxon tests, *n* = 836, *p <* 1 · 10^−16^ in both cases, Fig. 2G). Moreover, as expected for a differentiation trajectory from iPSCs to neural progenitors, the module eigengenes [4] of pluripotency markers were downregulated towards later pseudotime stages, and their target genes were enriched for pluripotency-related pathways such as *Transcriptional regulation of pluripotent stem cells* (Fig. 2K,L). In contrast, the module eigengenes of early neural markers showed increasing expression over pseudotime and their target genes were enriched for pathways such as *Formation of the anterior neural plate* (Fig. 2K,L).

Next, we hypothesized that a good module should also be enriched for direct targets of the central regulator. We first tested this hypothesis indirectly by examining the enrichment of the regulator’s binding motifs within cis-regulatory elements of its target genes. We indeed observed the expected enrichment in the pruned modules compared to the random and initial modules (paired Wilcoxon tests, *n* = 836, *p <* 1 · 10^−16^ in both cases, Fig. 2H, Supplementary Fig. S3D). We then sought more direct evidence by leveraging existing ChIP-seq data for POU5F1 [27]. Consistent with our hypothesis, the fraction of genes with at least one associated POU5F1 ChIP-seq peak was significantly higher in the POU5F1 module than among other network genes (Fisher’s exact test, odds ratio = 5.72, *p* = 5 · 10^−6^, Fig. 2I).

The above analyses demonstrate that combining network metrics with our novel pruning approach produces meaningful modules. But are these modules actually superior to the commonly used strategy of simply selecting the 50 strongest targets of each regulator [5, 44]? We can evaluate module quality using the phylogenetic signal strength and the difference in preservation between actual and random modules (see next section). We found that the dynamic pruning approach outperformed the top-50 strategy in both aspects (Supplementary Fig. S4A-H).

Taken together, these results indicate that the CroCoNet consensus modules capture biologically relevant information and provide a better balance between false positives and false negatives than fixed-size approaches, leading to a higher signal-to-noise ratio for cross-species comparisons.

### Preservation scores to compare module topologies

After identifying consensus modules, we defined a metric to compare module topologies across replicates and species. We relied on the joint module assignment derived from the consensus network but compared connection patterns directly between the replicate-wise networks. To define suitable metrics, we incorporated into CroCoNet two preservation statistics introduced by Langfelder *et al.* as part of the WGCNA workflow. The correlation of adjacencies (*cor.adj*) captures fine-grained topology at the level of individual edges, while the correlation of intramodular connectivities (*cor.kIM*) reflects higher-level topology based on intramodular connectivities — gene-level summaries of the underlying adjacencies (Supplementary Fig. S5A). To assess the suitability of these metrics for cross-species scRNA-seq analysis, we compared their values for actual versus random modules and examined the strength of the phylogenetic signal.

Both statistics were higher for the actual modules compared to the random modules (Fig. 3A; Supplementary Fig. S5B). Moreover, the average preservation followed the expected pattern: within-species comparisons yielded the highest scores, followed by human–gorilla comparisons, and then great ape (human & gorilla) versus cynomolgus macaque comparisons (Fig. 3A; Supplementary Fig. S5B).

**Figure 3.**
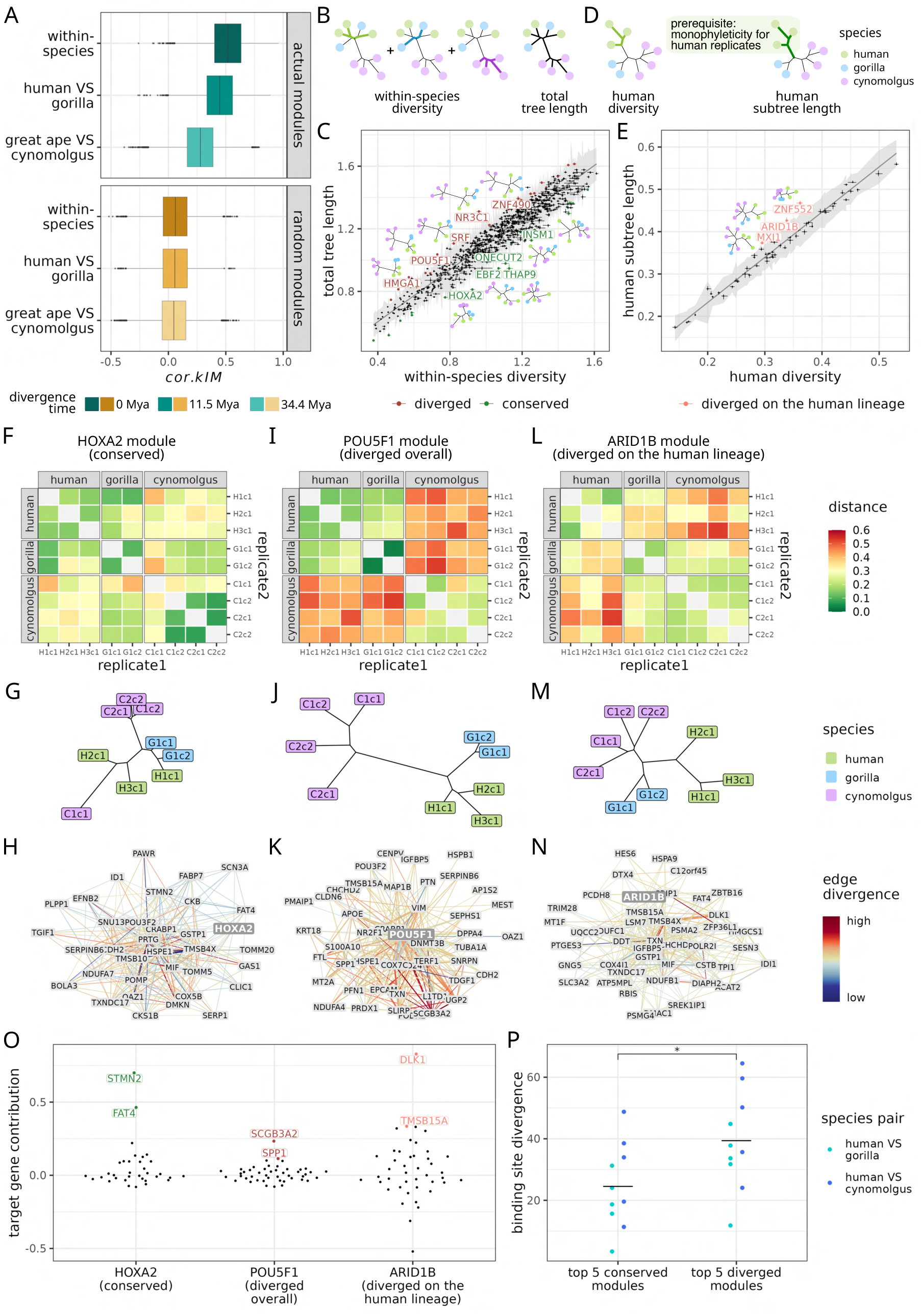
Quantifying cross-species conservation of gene regulatory networks. **A**) Distribution of *cor.kIM* for the actual and corresponding random modules, split by divergence time of the replicates compared. **B**) Total tree length and within-species diversity. **C**) Quantification of overall module divergence. A linear model was fitted between the total tree lengths and within-species diversities across all modules, and the 95% prediction interval of the regression line (depicted as a grey area) was calculated. Modules that fell above the upper bound or below the lower bound of the prediction interval were considered diverged and conserved, respectively. The 5 most conserved and 5 most diverged modules are labeled and displayed using the tree representations. **D**) Human diversity and human subtree length. The human subtree length is only defined if the tree is robustly monophyletic for the human replicates. **E**) Quantification of module divergence on the human lineage. A linear model was fitted between the human subtree lengths and human diversities across all human-monophyletic modules, and the 95% prediction interval of the regression line (shaded in grey) was calculated. Modules that fell above the upper bound of the prediction interval were considered diverged (labeled and displayed using the tree representations). **F**) **I**) **L**) Distance matrices based on *cor.kIM* for the modules HOXA2 (conserved), POU5F1 (diverged overall) and ARID1B (diverged on the human lineage). **G**) **J**) **M**) Neighbor-joining trees for the HOXA2, POU5F1 and ARID1B modules. **H**) **K**) **N**) The 200 strongest connections of the HOXA2, POU5F1 and ARID1B modules. The edge thickness represents the consensus edge weights (scaled per module) and the edge color represents how different the mean edge weights are across the three species (−log_10_*F*). **O**) Target gene contributions for the HOXA2, POU5F1 and ARID1B modules calculated based on jackknifing. For conserved modules, top-contributing targets drive the conservation signal, whereas for diverged modules, they drive the divergence signal. **P**) Binding site divergence of the regulators associated with the five most conserved and five most diverged modules. The binding site divergence is based on scoring all annotated motifs of the regulators in the ATAC-seq peaks of their module member genes.

Although both metrics appeared suitable by these criteria, *cor.kIM* provided better discriminatory power than *cor.adj* for the neural differentiation dataset (Supplementary Fig. S5C–G). In contrast, in the brain dataset, *cor.adj* more effectively distinguished actual from random modules and better captured the expected phylogenetic relationships (Supplementary Fig. S1E–J). This likely reflects differences in signal-to-noise ratio between the datasets. The differentiation data contain relatively few cells along a continuous developmental trajectory, favoring the use of the robust *cor.kIM*, whereas the brain data comprise roughly twenty times more cells from distinct, fully differentiated cell types favoring the more fine-grained *cor.adj*. Because the signal-to-noise ratio of a dataset is generally unknown, we recommend using the functions provided in CroCoNet to identify the optimal preservation metric.

### Quantifying module conservation

It is important to recognize that standard module preservation metrics reflect all sources of variance — not only evolutionary divergence but also technical and environmental noise, which may differ substantially across modules. Thus, to meaningfully interpret preservation metrics as proxies for evolutionary conservation, we must first account for the contributions of technical and environmental variation (Fig. 3B-E).

To accomplish this, we first summarize the preservation scores as neighbor-joining trees (Fig. 1C). The tips of these trees represent the replicates and the branch lengths are proportional to differences in module topology between the replicates. Like the preservation scores themselves, these branch lengths are affected by biological and technical confounding factors. To establish a baseline for these confounding factors, we use the within-species diversity, which we measure as the total length of all within-species subtrees (Fig. 3B). We capture the expected contribution of the confounding factors to the total module variability by fitting a linear regression model between the total tree length and the within-species diversity across all modules (Fig. 3C). Modules in the lower-left corner yield short trees, suggesting robustness to technical and environmental noise. Modules in the upper-right corner produce trees with long branches and little phylogenetic signal (Fig. 1D). Indeed, random modules also fall into the upper-right corner, supporting the interpretation that these trees are dominated by technical and environmental noise. We therefore exclude modules that fall within the distribution of random modules (Supplementary Fig. S4F).

Finally, the residuals from the regression line reflect the variance of the total tree length that cannot be explained by within-species diversity. In biological terms, this component of variance captures the contribution of between-species differences to the overall topological variability of a module. Positive residuals correspond to a greater-than-expected contribution of between-species divergence to total tree length, and negative residuals to lower-than-expected contributions. In order to identify modules with significant deviations from the overall trend, we calculated the 95% prediction interval of the linear fit and considered modules outside this prediction interval as conserved or diverged (Fig. 3C, Supplementary Tables 5-6).

For the differentiation dataset, we identified 20 conserved and 24 diverged modules (Supplementary Fig. S6A-F). Among the most conserved modules, we found several expected regulators. For example, HOXA2 (Fig. 3F-H), like many other Hox genes, shows strong conservation in its sequence, regulatory landscape, and function across vertebrates [45–47]. In contrast, the most diverged module is associated with NR3C1, the gene encoding the glucocorticoid receptor, which has a largely conserved protein sequence as well [48], but shows notable differences in hippocampal expression between humans and rhesus macaques [49]. Interestingly, among the most diverged modules, we also find POU5F1 (OCT4), a pluripotency factor that is deemed to have high functional and sequence conservation [48, 50], and shows a very conserved expression pattern in our data (Fig. 3I-K, Fig. 4A).

**Figure 4.**
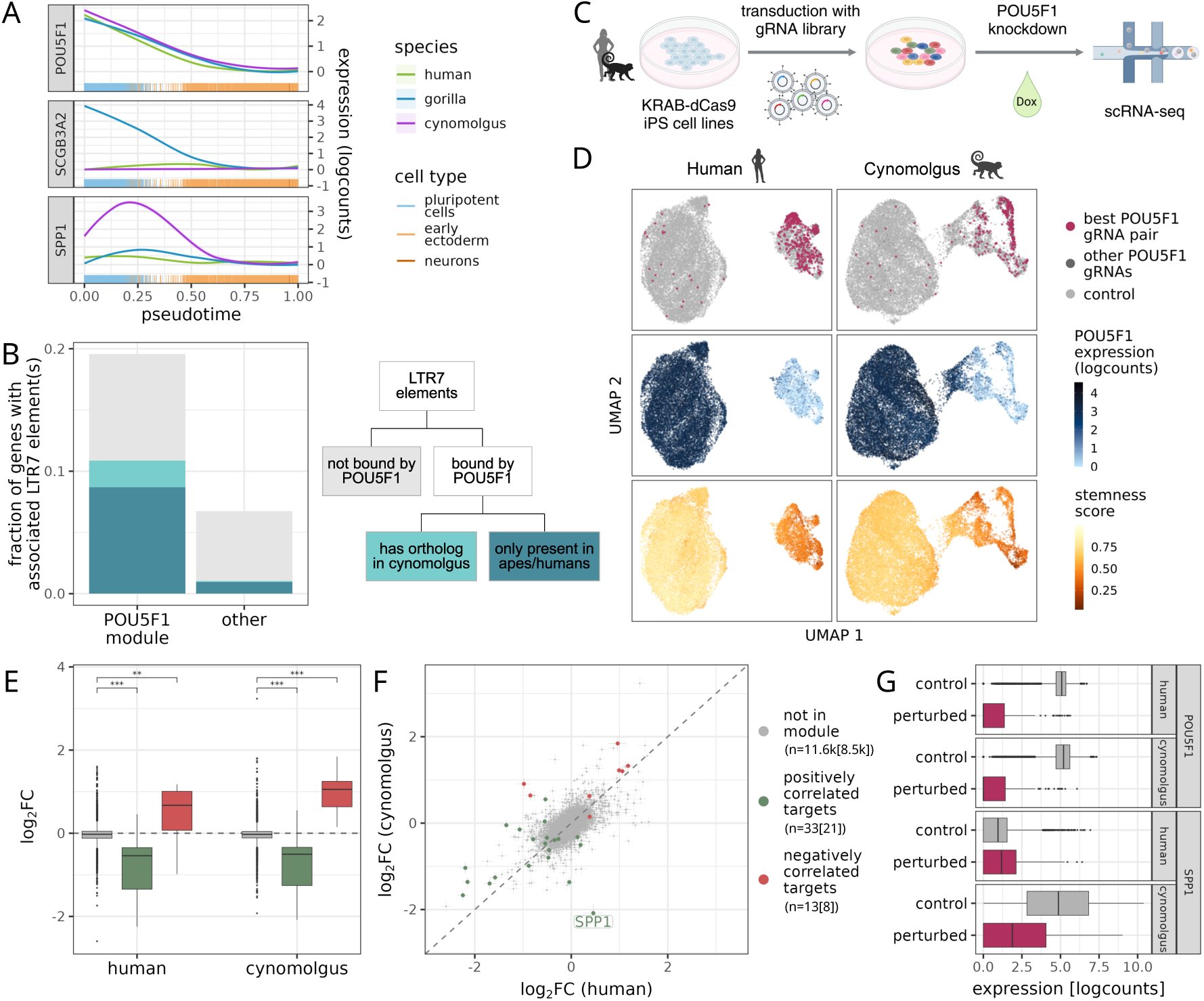
Additional evidence for the divergence of the POU5F1 module. **A**) Expression profiles of POU5F1, and its two most diverged targets, SCGB3A2 and SPP1. **B**) LTR7 enrichment near POU5F1 module member genes. Compared to all other network genes, in the vicinity of the POU5F1 module members we detected an enrichment of LTR7 elements (Fisher’s exact test, odds ratio = 3.37, *p* = 0.003), POU5F1-bound LTR7 elements (Fisher’s exact test, odds ratio = 11.6, *p* = 0.0001) and POU5F1-bound LTR7 elements that are only present in great apes or humans (Fisher’s exact test, odds ratio = 9.99, *p* = 0.001). **C**) Experimental setup of the single-cell CRISPRi screen. Two human and two cynomolgus macaque iPS cell lines with inducible dCas9-KRAB constructs were transduced with species-specific gRNA libraries containing POU5F1-targeting and control gRNAs. Five days after dCas9 induction with doxycycline, the gRNAs and transcriptome were profiled using scRNA-seq. **D**) UMAP representation of the POU5F1-perturbed and control cells. From top to bottom, cells are colored by gRNA identity, POU5F1 expression in logcounts, and stemness score. **E**) Log fold changes of positively correlated (in green) and negatively correlated (in red) CroCoNet POU5F1 module member genes as well as non-module genes (in grey) upon POU5F1 perturbation in human and cynomolgus. Asterisks indicate significance: * *p <* 0.05, ** *p <* 0.01, *** *p <* 0.001. **F**) Log fold changes upon POU5F1 perturbation in human and cynomolgus. Gene counts of positively and negatively correlated POU5F1 module member genes and non-module genes are indicated below the legend elements, with the number of genes also expressed in the CRISPRi screen shown in brackets. SPP1, the most differentially regulated gene in the CRISPRi screen, is labeled. **G**) Expression of POU5F1 and SPP1 in POU5F1-perturbed and control cells in human and cynomolgus macaque.

To identify impactful genes within diverged and conserved modules, we developed a metric to assess the contribution of each target gene to the overall module conservation score based on jackknifing (*c_i_*, Supplementary Fig. S7, see Methods). A positive score indicates that the presence of a target gene strengthens the signal of conservation or divergence, while a negative score means that the target gene weakens the signal. If *c_i_* ≥ 1, the categorization of the module as diverged or conserved was solely dependent on one gene. This was the case only for one module centered on the regulator PNRC2; for most modules, multiple genes contribute to the module conservation status (Fig. 3O, Supplementary Fig. S8, Supplementary Table 7). Most importantly, this metric should direct the researcher to regulator–target interactions that are most promising for further investigation.

### Lineage-specific divergence

The approach described above for identifying conserved and diverged modules provides a measure of overall conservation across the entire tree. However, because the longest lineages have the strongest influence, this conservation metric is dominated by the divergence between cynomolgus macaques and great apes in the neural differentiation dataset.

In many cases, however, the primary interest lies in lineage-specific selection, for which one would need to identify accelerated evolution along the lineage of interest. To address this, we need a well-defined lineage; for example, we define the human lineage as the branch leading from the most recent common ancestor (MRCA) of humans and gorillas to the human MRCA, using the macaque as an outgroup (Fig. 3D). A minimal requirement for this analysis is that the human replicates form a monophyletic group. Next, we quantify module conservation by contrasting the length of the human subtree, including the human lineage, to the human diversity (Fig. 3D). This allows us to assess module divergence specifically on the human lineage (Fig. 3E; Supplementary Table 5).

Following this approach, we identified three modules that are diverged on the human lineage (Supplementary Fig. S6G-I), including ARID1B, a chromatin remodeler with reduced expression in the human brain compared to the brains of other primates [51, 52] (Fig. 3L-N). Analogous to this human-centered analysis, an analysis focusing on the gorilla lineage identified 11 diverged modules, with CEBPB as the most diverged one (Supplementary Fig. S9, Supplementary Table 5).

### Interpreting topological divergence: cis versus trans contributions

Ultimately, a module’s conservation is determined by the sum total of cis- and trans-regulatory effects. On the one hand, if trans-effects are the primary drivers of module conservation, we expect the regulators of diverged modules to have either more diverged expression patterns or more diverged protein-coding sequences than the regulators of conserved modules. Indeed, for certain diverged modules such as NR3C1, the expression profiles of the regulators themselves differ markedly between species (Supplementary Fig. S8D). Overall, however, we do not find a significant difference in protein sequence or expression pattern divergence between the regulators of conserved and diverged modules (Supplementary Fig. S10A-B; Supplementary Table 5).

On the other hand, if the rewiring of a regulatory network occurs via changes in the cis-regulatory landscape, we expect to see a high divergence in the binding sites of the regulator within cis-regulatory elements (CREs) associated with its target genes. To obtain an estimate of binding site divergence, we identified putative CREs as ATAC-seq peaks within 20 kb from the transcription start site of each gene in iPSCs and NPCs, and then scored the overall binding potential of the regulator for each target gene and species (see Methods). Although such sequence-based binding predictions are far from reliable at the gene level, summarizing them across modules shows that the five most divergent modules indeed have higher binding site divergence than the five most conserved modules (Fig. 3P, Supplementary Fig. S10C, Supplementary Table 5). This finding suggests that cis-regulatory changes are important drivers of module divergence.

That said, cis- and trans-changes of a network are by no means mutually exclusive; both clearly contribute to network rewiring. It is noteworthy that CroCoNet – despite being solely based on co-expression – can detect module divergence arising from both cis- and trans-acting changes.

### Evidence for POU5F1 rewiring by LTR7 elements

The second most diverged module, POU5F1, is a potential example of cis-regulatory rewiring: the protein sequence is relatively conserved between human and cynomolgus macaque (99% amino acid identity) and 100% conserved between human and gorilla. Additionally, there is no evidence for differential regulation of POU5F1 itself (Fig. 4A). Moreover, based on our jackknife analysis, the rewiring does not hinge on a single target gene; instead, cis-regulatory changes must have affected many module members (Supplementary Fig. S7B, Fig. 3O). Regulatory changes are particularly evident when inspecting the expression trajectories of SCGB3A2 and SPP1, the two target genes that contribute the most to module divergence (Fig. 4A).

One proposed mechanism for network rewiring is the exaptation of transposable elements [53]. Consistent with this idea, in the human genome both SCGB3A2 and SPP1 are located near transposable elements from the LTR7 family, which has been shown to carry binding sites for multiple pluripotency factors, including POU5F1 [54–56]. To assess whether this extends beyond these two genes, we investigated the presence of LTR7 elements near the promoters of all POU5F1 module member genes in the human genome (Fig. 4B). Compared to all other genes, we detected a significant enrichment of LTR7 elements in the vicinity of POU5F1 module members (Fisher’s exact test, odds ratio = 3.37, *p* = 0.003), and an even stronger enrichment of LTR7 elements that are bound by POU5F1 in human embryonic stem cells based on ChIP-seq data (Fisher’s exact test, odds ratio = 11.6, *p* = 0.0001) [57]. Furthermore, many of the POU5F1-bound LTR7 elements near POU5F1 module member genes are great ape- or human-specific, including two LTR7 elements upstream of SPP1 and two upstream of SCGB3A2. Therefore, we hypothesize that these young LTR7 elements contributed to the rewiring of the POU5F1 module on the great ape lineage.

### Validation of POU5F1 module divergence by CRISPRi

To validate the CroCoNet approach for assigning co-expression modules and assessing their divergence, we perturbed POU5F1 in two human and two cynomolgus iPS cell lines using CRISPR interference (CRISPRi) (Fig. 4C, Supplementary Tables 1, 8-9). CRISPRi perturbations enable the identification of target genes for a given regulator through a simple differential expression analysis between perturbed and unperturbed cells. When performed across species, differential expression comparisons provide a measure of regulatory divergence by identifying differentially regulated genes, i.e., genes whose responses to the knockdown differ between species (see Methods).

Using scRNA-seq, we confirmed the efficiency of the perturbation: cells expressing a POU5F1-targeting gRNA showed reduced POU5F1 expression and a marked decrease in the stemness index, indicating successful repression of POU5F1 (Fig. 4D). Of the 46 genes in the POU5F1 CroCoNet module, 43 were detected as expressed in the CRISPRi dataset and were therefore available for assessing module membership. The three excluded genes are expressed either only in gorilla iPSCs (SCGB3A2) or only in NPCs (NR2F1 and POU3F2), and thus fell below the detection threshold in the CRISPRi cell lines. Strikingly, all 43 expressed module genes were identified as POU5F1 targets in the CRISPRi differential expression analysis, providing strong validation of the CroCoNet-derived consensus module (Fisher’s exact test, odds ratio = Inf, *p* = 2 · 10^−12^). Furthermore, the direction of regulation was highly consistent between the CroCoNet inference and the knockdown experiment: genes inferred to be repressed by POU5F1 were upregulated upon knockdown, and genes inferred to be activated were downregulated. In contrast, genes outside the module showed only minimal expression changes (Fig. 4E, Supplementary Table 10).

We also found that 28 of the 43 expressed module genes were significantly differentially regulated between the two species. This represents a significant enrichment compared to all other differentially expressed genes (Fisher’s exact test, odds ratio = 4.29, *p* = 3 · 10^−6^), confirming that the POU5F1 module is indeed diverged between human and cynomolgus iPSCs. The most strongly differentially regulated gene was SPP1 (Fig. 4F; log_2_ FC = −2.3, *p*_adj_ *<* 1 · 10^−16^), which was also identified as the second most diverged POU5F1 target in the CroCoNet analysis (the most diverged, SCGB3A2, could not be tested here; Fig. 3O). Consistent with the expression patterns observed in the neuronal differentiation dataset, SPP1 showed higher expression in unperturbed cynomolgus iPSCs than in unperturbed human iPSCs (Fig. 4G). Upon POU5F1 knockdown, SPP1 was slightly but significantly upregulated in human cells (log_2_ FC = 0.45, *p*_adj_ = 0.02), whereas it was significantly downregulated in cynomolgus cells (log_2_ FC = −1.9, *p*_adj_ *<* 1 · 10^−16^). This independently confirms that SPP1 is regulated differently by POU5F1 in humans and cynomolgus macaques.

Taken together, the CRISPRi perturbation experiment validates both the module membership and the regulatory divergence of the POU5F1 module across primates. This example demonstrates that CroCoNet recovers biologically meaningful cross-species differences in network wiring.

## Discussion

CroCoNet, the framework we introduce, is designed to analyze GRN divergence among closely related species, where cis-regulatory rewiring or expression changes of transcription factors are the main driving forces.

The biggest issue with evolutionary comparisons of GRN architectures is the low power to detect true connections. This is particularly problematic for single-cell data from closely related species, where a low signal-to-noise ratio is expected [23]. In order to alleviate this issue in a non-comparative setting, it has been suggested to integrate complementary data such as transcription factor binding [5] and information from protein–protein interactions [58]. Such other data types are still rare for non-model organisms, and thus integrating this kind of information in a cross-species comparison bears the risk of introducing a species bias. Thus, GRN comparisons across species have so far mainly been based on co-expression networks, and due to the power issues, the focus has been on conserved modules [3].

In CroCoNet, we tackle the issue of cross-species GRN comparison by 1) optimizing the identification of the consensus modules using a dynamic pruning approach, 2) adapting robust preservation statistics, and 3) using biological replicates to disentangle phylogenetic network divergence from within-species diversity, including environmental and technical variance, to provide an unbiased divergence score.

Because transcriptional regulation is often cooperative, we allow genes to belong to multiple modules rather than enforcing mutually exclusive membership. This choice is supported by the observation that modules whose central regulators interact tend to share more genes (Fig. 2J). Another critical point is striking the right balance between sensitivity and specificity in defining module membership. In practice, overly inclusive modules risk accumulating false positives, which in turn weakens any signal of conservation. To manage this trade-off, our dynamic pruning algorithm evaluates each gene’s adjacency to the central regulator as well as its connectivity to other module members. This refinement leads to more biologically coherent modules: the pruned sets display a stronger phylogenetic signal than an alternative approach that simply retains the 50 most connected genes (Supplementary Fig. S3D).

Beyond the qualitative module detection, the quantitative evaluation of module topology also critically depends on balancing sensitivity and specificity. In CroCoNet, we adapted two types of correlation-based preservation statistics that were suggested by Langfelder et al. [7]. The first one, *cor.adj*, compares the edges of a module, while the second one, *cor.kIM*, provides a higher-level summary by comparing the connectivities per gene (Supplementary Fig. S4A). In the differentiation dataset, *cor.kIM* outperforms *cor.adj*, showing a stronger phylogenetic signal (Supplementary Fig. S4B-G), while for the brain dataset *cor.adj* performs better (Supplementary Fig. S1E-G). We interpret this as reflecting the different signal-to-noise levels across the datasets: regulatory interactions inferred from the differentiation dataset are comparatively noisy, favoring the more robust summary statistic *cor.kIM*, whereas in the brain dataset, subtler — but still meaningful — differences are detectable only with the more fine-grained, edge-level measure *cor.adj*.

The preservation statistics and the resulting neighbor-joining trees reflect not only phylogenetic divergence but also within-species diversity, including variation due to environmental and technical factors. Here, we are mainly interested in the phylogenetic signal and treat the within-species variation as a nuisance parameter when quantifying module divergence. Modules with large total tree lengths are dominated by the nuisance parameter and their tree characteristics resemble those of random modules, while modules with very small overall tree sizes are robust (Supplementary Fig. S3F). In order to rank the modules according to their level of between-species divergence, we assume that the contribution of divergence to the total tree length is approximately constant across modules, and we look for deviations from this expectation. A larger contribution indicates accelerated evolution, while a smaller contribution indicates higher conservation. The logic behind this approach is in part analogous to a McDonald-Kreitman test, wherein the ratio of non-synonymous (=functional) to synonymous (=neutral) divergence is contrasted with that of diversity [20]. The main difference is that we have no measure of neutral evolution, and thus cannot quantify the amount of selection. Therefore, our scores can only be interpreted relative to all other modules. Nevertheless, the logic has been successfully applied to evaluate the evolution of gene expression levels [59–62], and hence is also valid in a network context, allowing us to identify diverged and conserved modules.

CroCoNet also provides tools to investigate these interesting modules in more depth. To begin with, we calculate a metric for the contribution of an individual target gene to the overall module divergence (Fig. 3O). Additionally, we provide functions to compare the expression trajectories of single genes as well as module eigengenes (Fig. 4A). These comparisons are particularly relevant when trying to distinguish cis- from trans-effects: if a module is diverged due to changes in the central regulator’s expression pattern, then that regulator should be detected as differentially expressed. This is, for example, the case for the most diverged module in the differentiation dataset, NR3C1 (Supplementary Fig. S8D). In contrast, POU5F1 (OCT4), the central regulator of the second most diverged module, has a very conserved expression trajectory (Fig. 4A), showing that CroCoNet is also equipped to detect modules that are diverged due to cis-regulatory rewiring.

Kunarso et al. (2010) [54] showed that although POU5F1 is a core pluripotency factor in both human and mouse, its genomic binding locations are largely non-orthologous, with transposable elements (TEs) contributing roughly one quarter of the binding sites in each species. Yet, only a subset of these TE-derived “proto-enhancers” exerts detectable regulatory effects [63, 64], leaving open the question of whether such cis-regulatory turnover meaningfully reshapes the gene regulatory network. Here, we show that this turnover indeed corresponds to measurable changes in the topology of the POU5F1 network. CroCoNet is well suited to this analysis, because it directly compares GRN topologies across species, allowing us to assess the extent to which cis-regulatory rewiring translates into network-level rewiring. What remains unclear is whether these network differences reflect adaptive divergence in early development, or instead represent developmental system drift, where changes in regulatory architecture do not alter core functional outcomes [65].

## Methods

### Experimental procedures and data processing

To generate the neural differentiation dataset, three human, two gorilla and four cynomolgus macaque iPS cell lines [66–70] (Supplementary Table 1) were differentiated into NPCs over the course of nine days using dual-SMAD inhibition [71, 72]. On days 0, 1, 3, 5, 7 and 9, cells were sampled and profiled using mcSCRB-seq [73].

Reads were mapped to hg38, gorGor6 and macFas6 using zUMIs [74]. For the human genome, we used the GENCODE gene build (release 32). For the gorilla and cynomolgus macaque, we transferred the human gene build to the corresponding genomes using Liftoff [25]. We removed low-quality cells and unexpected cell types (mainly neural crest), as well as genes with unsuccessful liftoff, low expression, and unwanted gene types (see Supplementary Methods). We normalized counts using scran [75] and annotated cell types using SingleR [76] with a human embryoid body dataset as the reference [77]. After batch effect removal [78], we inferred pseudotime using SCORPIUS [79].

For the brain dataset profiling the middle temporal gyrus of five primate species [22], we downloaded the filtered count matrices with the attached metadata from https://data.nemoarchive.org/publicationrelease/Great_Ape_MTG_Analysis/. We only kept 10x data and removed the donor H18.30.001 due to its atypical cell type composition as well as the donors H200.1023 and C19.32.006 due to their low cell numbers. This left us with four human, six chimpanzee, four gorilla, three rhesus and three marmoset donors (Supplementary Fig. S5A-B, Supplementary Table 2).

### Network inference

In case of the differentiation dataset, we normalized counts per replicate using randomized quantile residual transformations prior to network inference [80]. Based on these normalized counts, we inferred networks per replicate with GRNBoost2 [16, 81], specifying all genes in the filtered count matrix as potential regulators. We ran the algorithm 10 times for each replicate with 10 different seeds, then averaged edge weights across the 10 runs and 2 possible directions. Edges that reached the detection limit in none or only one of these 20 network versions were assigned a weight of 0. This ensured that, despite GRNBoost2 not providing importance scores for every possible edge, the resulting networks were still fully connected.

In case of the brain dataset, all donors were subsampled to the same number of cells per cell type, resulting in 9,456 cells per donor (Supplementary Fig. S5B). We created ten independent subsamples with this cell number and cell type composition per donor and performed ten independent network reconstructions to retain as much information from the data as possible. For each network reconstruction, we calculated Spearman’s correlations between all gene pairs based on the normalized expression matrices across all cell types. The network reconstructions of the different subsamples were then averaged to obtain a single network per donor. The correlation coefficients were transformed into unsigned adjacencies by taking their absolute values.

### Consensus network

We integrated the per-replicate networks into a consensus network in a phylogeny-aware manner, building on the concepts of Phylogenetic Generalized Least Squares (PGLS) [82, 83]. Based on primate divergence time estimates [84], we calculated a phylogenetic similarity matrix (*S*) across all replicates:

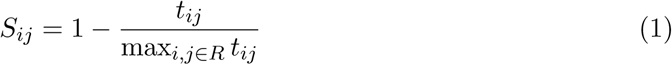

where *R* is the set of replicates for which the consensus is calculated, and *t_ij_*is the divergence time of the two species replicates *i* and *j* belong to.

Using this similarity matrix, the weights of the replicates (*w_i_*) were calculated as:

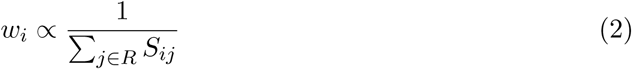

and were normalized to sum to 1. Finally, the consensus adjacency of an edge (*a_consensus_*) was calculated as the weighted mean of the adjacencies in each replicate (*a_i_*):

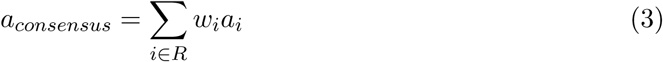

### Module assignment

As central regulators for the module assignment, we selected genes that had at least one annotated binding motif in the JASPAR [18] or IMAGE [19] databases and had a variance above noise level in the data based on variance decomposition [75]. Applying these criteria yielded 836 regulators for the differentiation dataset and 499 regulators for the brain dataset.

Modules associated with these regulators were created and pruned based on the consensus network. Specifically, we used two metrics: 1) the regulator–target adjacency (*a*_regulator_), which is the adjacency between the regulator and a gene, and 2) the intramodular connectivity (*kIM*), which is the sum of a gene’s adjacencies to all other genes within the same module [7]:

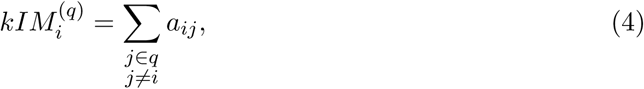

where *q* is the set of genes that belong to a module.

We assigned an initial module to each regulator by selecting its top 4,000 target genes based on *a*_regulator_. Next, we pruned the modules using an iterative two-step process: 1) we calculated the cumulative sum curve based on *a*_regulator_ per module, identified the knee point of the curve [85], then kept the targets with a higher *a*_regulator_ than the knee point, and 2) we repeated the knee-point–based selection for *kIM*. Pruning steps 1) and 2) were repeated until the module sizes became as small as possible without any of them falling below 20 (Supplementary Tables 3-4). For the differentiation dataset, pruned module sizes ranged between 25 and 63, with a median of 45, while for the brain dataset, module sizes ranged between 29 and 67, with a median of 51.

Corresponding to each module, we also created a random module of the same size by randomly drawing genes from the network (without replacement).

### Inferring binding potential and binding site divergence of the regulators

We validated the membership and divergence of the modules in the differentiation dataset based on binding site predictions of the central regulators. To this end, ATAC-seq data were collected from two human, two gorilla and three cynomolgus iPSC lines as well as from two human and two cynomolgus NPC lines [67–70] (Supplementary Table 1) using the Omni-ATAC protocol [86]. Reads were mapped to the same reference genomes as for the scRNA-seq data using BWA-MEM2 [87]. Peaks were called jointly for all replicates per cell type and species using genrich [88]. Peaks in gorilla NPCs were inferred by LiftOver [89] of the human NPC peaks to gorGor6.

Active transcription start sites (TSS) were identified in human, gorilla and cynomolgus iPSCs and NPCs using the GENCODE or Liftoff annotations, long read RNA-seq data and ATAC-seq data (see Supplementary Methods). For each cell type and species, ATAC-seq peaks were associated with genes based on their distance to TSSs [69], allowing a maximum distance of 20 kb. We scored the binding motifs of each central regulator [19, 90] in all peaks associated with the genes in their initial, pruned and random modules using Cluster-Buster [91]. After collapsing redundant motif hits, we calculated the binding potential of a regulator per gene and species by summing the motif scores across all associated peaks. For module validation, the binding potential was further summarized per module using the median.

To investigate cross-species divergence in binding potential, we calculated the absolute differences in binding potential between all species pairs per gene and summarized these differences again as median per species pair and module (Supplementary Table 5). To test whether regulators with conserved and diverged network modules differ in terms of their binding site divergence, we fit the following linear model across the human-gorilla and human-cynomolgus comparisons of conserved and diverged CroCoNet candidates:

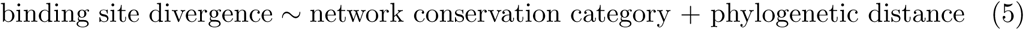

When contrasting all conserved to all diverged modules, we detected no significant difference in binding site divergence (*β* = 3.48, *p* = 0.47), but when contrasting the top five modules from each category, we detected a significantly higher binding site divergence for the regulators of diverged modules (*β* = 14.9, *p* = 0.02).

### Module preservation, tree reconstruction and module filtering

We quantified module preservation by calculating the preservation statistics *cor.adj* (correlation of adjacencies) and *cor.kIM* (correlation of intramodular connectivities) [7] for each consensus module between all possible pairs of replicates.

We converted the preservation scores of the chosen statistic (*pres*) into distances (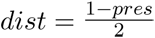) to construct a distance matrix and infer a neighbor-joining tree per module [92]. Based on these trees, we calculated overall tree characteristics (the total tree length and within-species diversity, Fig. 3B) as well as lineage-specific tree characteristics (subtree length and diversity of the species of interest, Fig. 3D).

To assess the uncertainty these metrics, we used jackknifing, meaning that we removed each member gene of a module and re-calculated all preservation scores and tree-based statistics. We used the median and its 95% confidence interval across the jackknifed versions to estimate a statistic of interest for the module as a whole.

We used the overall tree characteristics to remove uninformative modules. For each module, we estimated the probability of originating from the distribution of random modules (*p*_random_) and from the distribution of actual modules (*p*_actual_), using the probability density functions of the two bivariate normal distributions. If

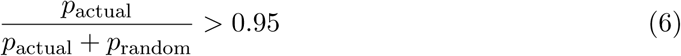

was not met, we excluded the module. This led to the removal of 10 modules in case of the differentiation dataset and 11 modules in case of the brain dataset.

### Quantification of cross-species conservation

To characterize the overall divergence of the modules, we fitted a weighted linear model with the total tree length (*L_t_*) as dependent, and the within-species diversity (*L_w_*) as independent variable (*L_t_* = *βL_w_* + *c* + *ε*). Specifically, for each module we used the median total tree length and median within-species diversity across its jackknifed versions and weighted the data point inversely proportional to the jackknife variance estimate of the total tree length. We regarded the residuals of each module *m* (*ε_m_*) from this linear model as measures of cross-species module divergence. To gain information about the significance of these divergence scores, we calculated the 95% prediction interval of the regression line. We considered a module significantly diverged if it had a higher total tree length than the upper boundary of the prediction interval, while we considered a module significantly conserved if it had a lower total tree length than the lower boundary of the prediction interval (Supplementary Tables 5-6). In case of For the differentiation dataset, we detected 20 conserved and 24 diverged modules (Fig. 3C), while for the brain dataset, we detected 10 conserved and 15 diverged modules (Supplementary Fig. S5J).

To estimate the lineage-specific divergence of a module, the module tree was required to be monophyletic for the species of interest across all jackknife versions. Analogous to the analysis of overall divergence, we fitted a weighted linear model, this time using the species-specific subtree length instead of the total tree length and species-specific diversity instead of the overall within-species diversity. Note that due to the filtering for monophyleticity, we lose the most conserved modules and thus cannot interpret modules below the lower boundary of the prediction interval as conserved.

### Target gene contribution

To identify the most conserved and diverged target genes within a module, we used the results of jackknifing. We relied on the linear model that was fit across all modules between the total tree length and within-species diversity (for overall divergence), or between the subtree length and diversity of a species (for lineage-specific divergence). For each jackknifed module version created by removing a gene *i*, we re-calculated the total tree length (*L_t,i_*) and within-species diversity (*L_w,i_*), and, based on these, we also obtained the deviation from the original regression line (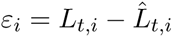, Supplementary Table 7). Using the residual of the original module (*ε_m_*), we defined the target gene contribution score (*c_i_*) as follows:

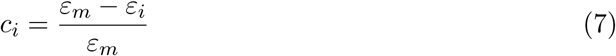

### POU5F1 perturbations using CRISPRi

We perturbed POU5F1 in human and cynomolgus inducible KRAB-dCas9 iPS cell lines [70] (Supplementary Table 1) as part of three pooled single-cell CRISPRi screens. We designed gRNAs targeting POU5F1 (Supplementary Table 8) as well as non-targeting control gRNAs (Supplementary Table 9). Details of the gRNA design, experimental procedures and initial data processing are described in Edenhofer *et al.* (2024) [70] and in the Supplementary Methods.

For the validation of the CroCoNet POU5F1 module in the differentiation dataset, we selected one gRNA per species with high and comparable knockdown efficiencies (human: GACAGAACTCATACGGCGGG, % KD = 83.9%; cynomolgus: GAAGCCAGGTGTCC-CGCCAT, % KD = 84.8%). To further ensure comparability, we kept only batches shared across the two gRNAs and downsampled to an equal number of cells per condition and species, resulting in 288 POU5F1-perturbed and 288 unperturbed control cells in both human and cynomolgus.

Next, we fitted a linear mixed model using dream [93] with the design:

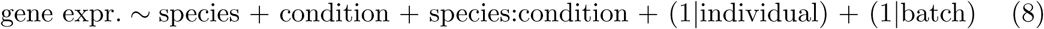

We tested three contrasts: 1) perturbed vs. control in human cells, 2) perturbed vs. control in cynomolgus cells, and 3) the interaction between species and condition (i.e. evaluating whether the perturbed–control difference is significantly larger in one of the species). We regarded a downstream gene as significantly differentially expressed (DE) within a species or significantly differentially regulated (DR) across the species if the corresponding contrast had an adjusted p-value *<*0.05 (Benjamini-Hochberg FDR correction). Out of the 10,207 genes tested, 3,871 were DE in human, 3,471 were DE in cynomolgus and 1,720 were DR across the two species, while out of the 43 POU5F1 module member genes tested, 39 were DE in human, 37 were DE in cynomolgus and 28 were DR across the two species (Supplementary Table 10).

## Abbreviations

ATAC-seq: assay for transposase-accessible chromatin using sequencing
CCDS: canonical coding sequence
CDS: coding sequence
ChIP-seq: chromatin immunoprecipitation sequencing
*cor.adj*: correlation of adjacencies
*cor.kIM*: correlation of intramodular connectivities
CRE: cis-regulatory element
CRISPRi: clustered regularly interspaced short palindromic repeats interference
DE: differential expression/differentially expressed
DR: differential regulation/differentially regulated
FDR: false discovery rate
GEX: gene expression
GRN: gene regulatory network
iPSC: induced pluripotent stem cell
kIM: intramodular connectivity
MOI: multiplicity of infection
MRCA: most recent common ancestor
NPC: neural precursor cell
PCBC: Progenitor Cell Biology Consortium
PGLS: Phylogenetic Generalized Least Squares
scRNA-seq: single-cell RNA sequencing
SNP: single nucleotide polymorphisms
TE: transposable element
TR: transcriptional regulator
TSS: transcription start site
UMI: unique molecular identifier
UTR: untranslated region

## 1 Declarations

### Consent for publication

Not applicable.

### Availability of data and materials

The CroCoNet R package is available at https://github.com/Hellmann-Lab/CroCoNet, with detailed documentation and a step-by-step tutorial at https://hellmann-lab.github.io/CroCoNet/. The code for all analyses in this manuscript is available at https://github.com/ Hellmann-Lab/CroCoNet-analyses, data files are deposited in the linked Zenodo archive. The neural differentiation scRNA-seq data was deposited to ArrayExpress under the accession number E-MTAB-15695. The ATAC-seq data was deposited to ArrayExpress under accession numbers E-MTAB-13373 and E-MTAB-15654. The single-cell CRISPRi data was deposited to GEO under the accession number GSE298717

### Author’s contributions

IH and AT conceptualized this study. IH acquired funding. AT implemented the CroCoNet R package. JG performed the neural differentiation scRNA-seq experiment. AT processed the scRNA-seq data, performed the network inference, applied the CroCoNet workflow to the example data, run the additional analyses for validation and visualized the results. VS, AT and ZK did the ATAC-seq analysis and motif scoring. VS analyzed the POU5F1 ChIP-seq data. FCE performed the single-cell CRISPRi experiments. BV, ZK and PJ provided guidance and scripts for the analysis. IH supervised the project. IH and AT wrote the original draft. All authors reviewed and edited the manuscript.

## Acknowledgements

We would like to thank Wolfgang Enard for his support and feedback throughout the project. We are grateful to Paulina Spurk and Daniel Richter for the generation and preprocessing of the ATAC-seq dataset. We acknowledge the Core Facility Flow Cytometry at the Biomedical Center, LMU Munich for providing equipment and expertise. We thank the Laboratory for Functional Genome Analysis (LAFUGA) at the Gene Center, LMU Munich, in particular to Stefan Krebs and Helmut Blum, as well as the NGS Competence Center Tübingen (NCCT) for their sequencing services. We are grateful to all members of the Hellmann-Enard group for the discussions and valuable input.

## Supplementary Information

### Supplementary figures

**Supplementary Figure S1.**
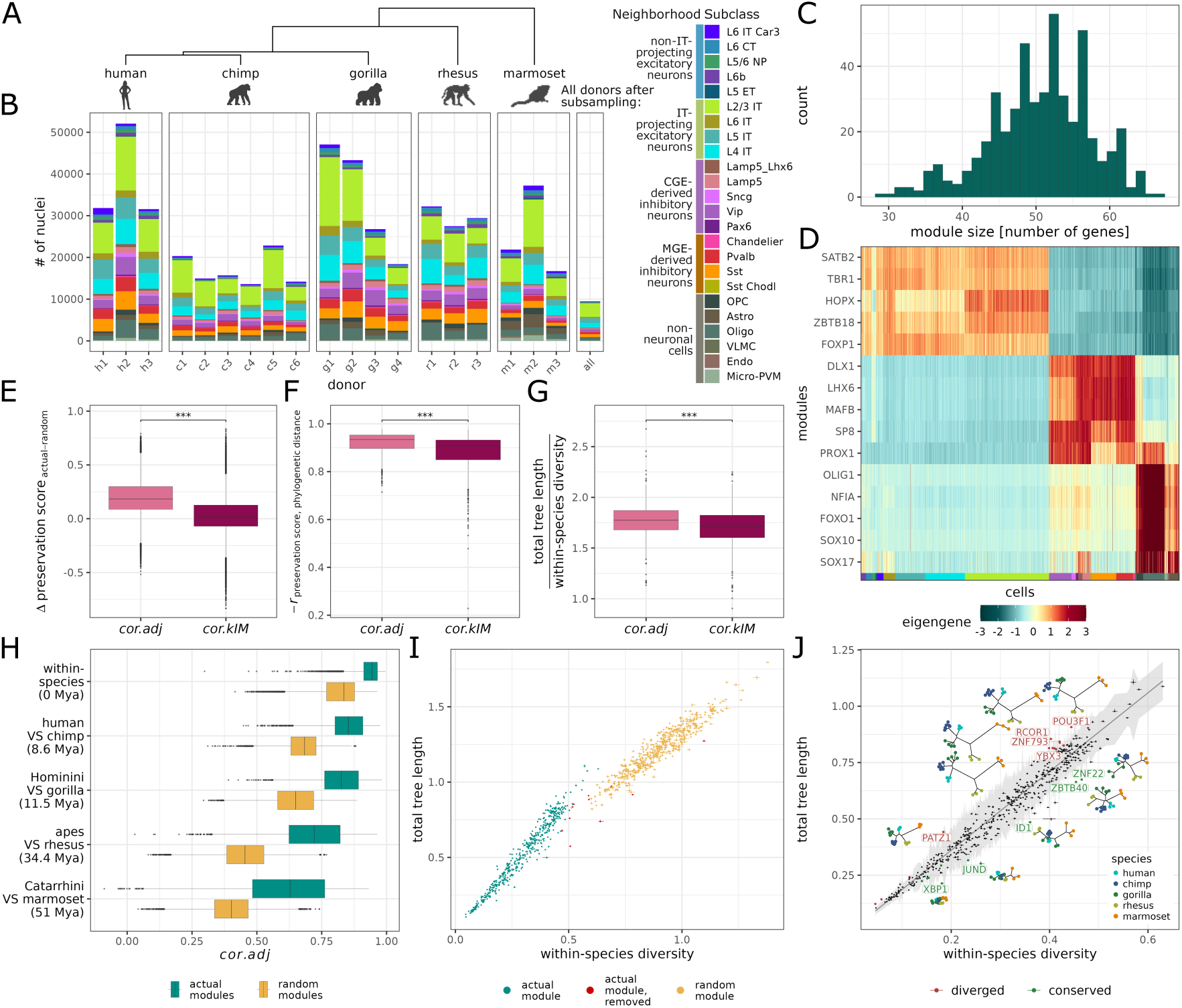
CroCoNet analysis of a single-nucleus RNA-seq dataset profiling the middle temporal gyrus of five primate species. **A**) Phylogenetic tree of the five species. **B**) Number of nuclei and cell type composition for each of the 19 donors after filtering. The last column shows the nuclei number and cell type composition each donor was subsampled to for network analysis. **C**) Distribution of module sizes after pruning. **D**) Module eigengenes of five excitatory neuronal, five inhibitory neuronal and five non-neuronal markers. The eigengene calculation was based on the positively correlated targets. **E**) Difference in the *cor.kIM* /*cor.adj* scores between each actual and the corresponding random module. **F**) Inverted Pearson’s correlation (−*r*) between the *cor.kIM* /*cor.adj* scores and phylogenetic distance. **G**) Ratios of the total tree lengths and within-species diversities in the tree representations of the pruned modules, reconstructed based on the *cor.kIM* and *cor.adj* scores. **H**) Distribution of the *cor.adj* scores for the pruned and corresponding random modules, split by divergence time of the replicates compared. **I**) Total tree lengths and within-species diversities of the pruned and corresponding random modules, based on the trees reconstructed using the *cor.adj* scores. Eleven actual modules (in red) were not distinct enough from the random modules and were therefore excluded from the module conservation analysis. **J**) Quantification of overall module conservation. A weighted linear model was fitted between the total tree lengths and within-species diversities across all modules, and the 95% prediction interval of the regression line (shaded in grey) was calculated. Modules that fell above the upper bound or below the lower bound of the prediction interval were considered diverged and conserved, respectively. The five most conserved and five most diverged modules are labeled and displayed using the tree representations.

**Supplementary Figure S2.**
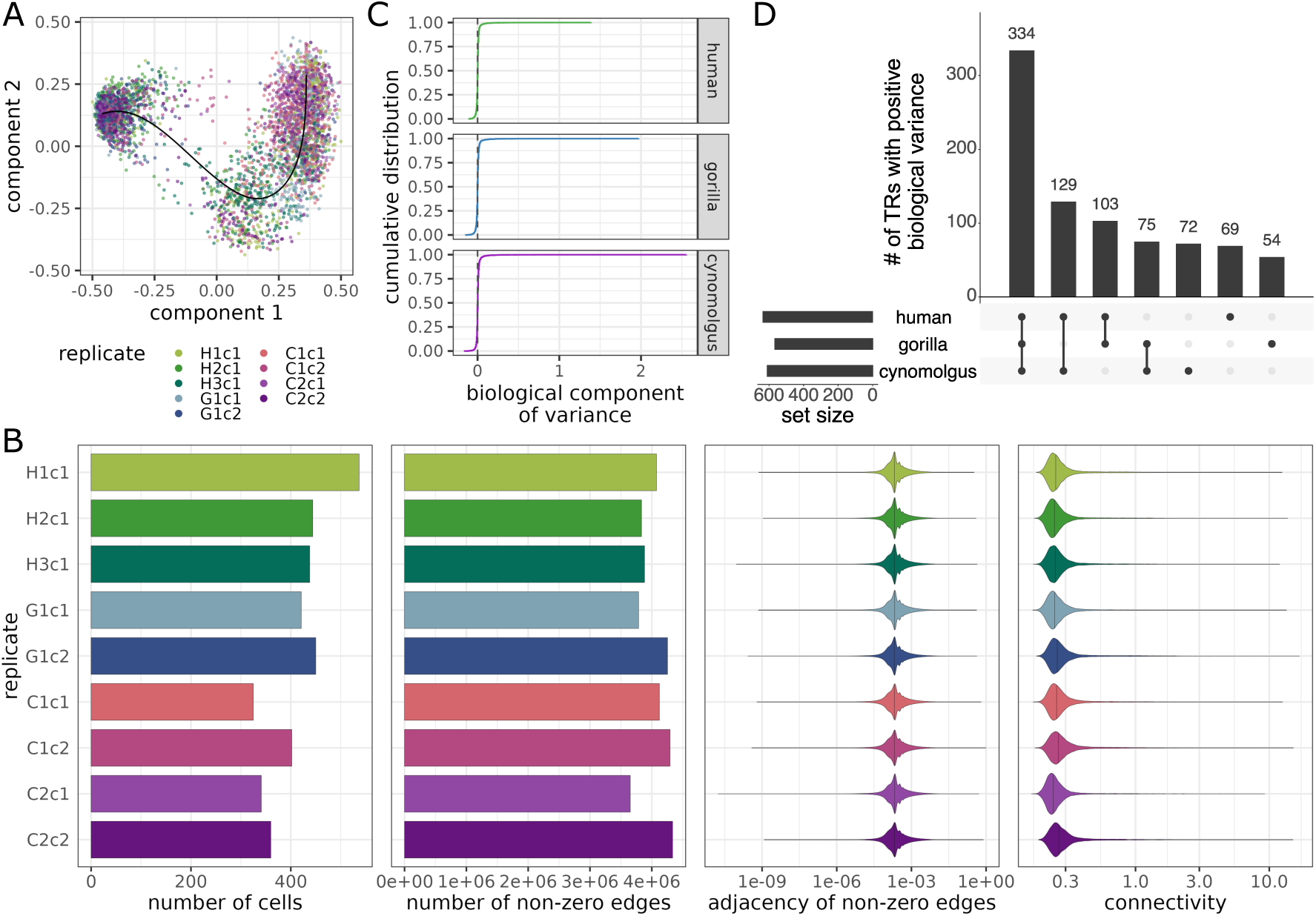
Main characteristics of the replicate-wise networks and the construction of the consensus network. **A**) Pseudotime trajectory jointly inferred for all species, with cells colored by replicate. **B**) The number of cells that the network inference was based on, the number of non-zero edges, the distribution of adjacencies across all non-zero edges, and the distribution of connectivities across all genes for each replicate-wise network. **C**) Selection of relevant transcriptional regulators based on biological variance. The variance of the normalized gene counts was decomposed into technical and biological components, and TRs that had a positive biological variance in any of the species (cutoff indicated by the dashed line) were used as central regulators for the module assignment (*n* = 836). **D**) TRs with a positive biological variance shared between different subsets of the species.

**Supplementary Figure S3.**
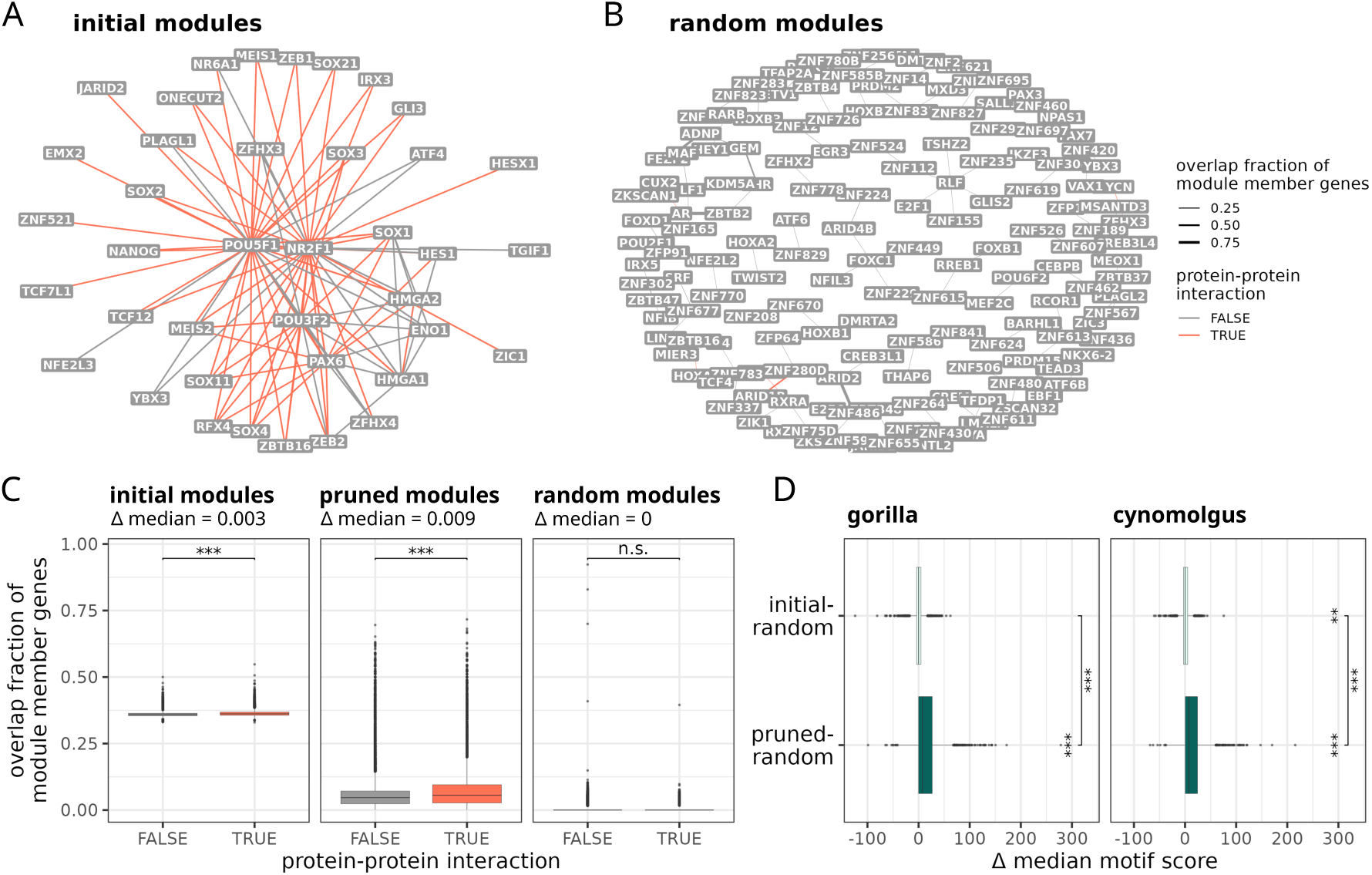
Validation of the modules based on motif enrichment and target gene overlap. **A-B**) Overlaps of the initial and random modules and protein–protein interactions among their regulators. Nodes represent the initial/random modules associated with the regulators on the labels, and edge weights represent the overlap fraction of module members. Only the 100 highest overlaps are shown. Red edges indicate that the regulators interact based on STRINGdb [28]. **C**) Overlap of module member genes for interacting versus non-interacting regulators. Interactions were defined based on STRINGdb [28]. The overlap between two pruned or initial modules was significantly higher if the corresponding regulators were interaction partners (Wilcoxon test, *n*_1_ = 17, 670, *n*_2_ = 331, 360, *p <* 1 · 10*^−^*^16^ in both cases), while the same was not observed for random modules (*p* = 0.45). The difference in the median overlaps (shown in the subtitles) was the highest for the pruned modules. **D**) Each target gene was scored for binding motifs of the module’s central regulator in the gorilla and cynomolgus genomes (see Methods). The plot shows how the module-level summaries of these motif scores differ in the initial and pruned modules relative to the random modules for each species. **C–D**) Asterisks indicate significance: * *p <* 0.05, ** *p <* 0.01, *** *p <* 0.001.

**Supplementary Figure S4.**
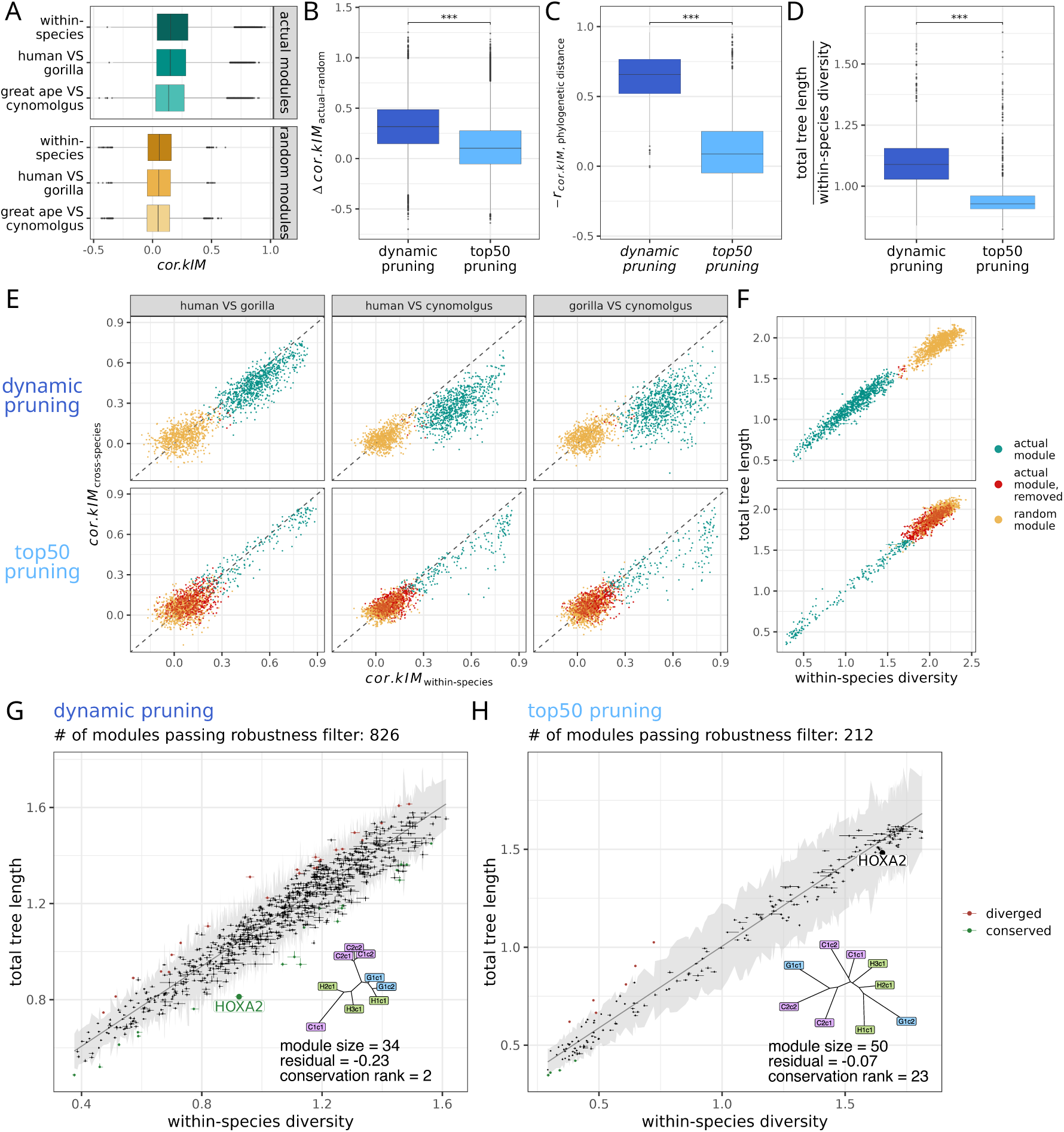
Comparison of the dynamic and top50 pruning approaches. **A**) Distribution of *cor.kIM* for modules pruned using the top50 approach and the corresponding random modules, split by divergence time of the replicates compared. **B**) Difference in the *cor.kIM* scores between each actual and the corresponding random module, for the dynamic and top50 pruning. **C**) Inverted Pearson’s correlation (−*r*) between *cor.kIM* and phylogenetic distance for the dynamic and top50 pruning. **D**) Ratios of the total tree lengths and within-species diversities in the tree representations of the pruned modules created by dynamic and top50 pruning. **E**) Within-species and cross-species *cor.kIM* scores of the pruned and corresponding random modules, split by pruning approach (top50 or dynamic) and species pair. Actual modules in red were not distinct enough from the random modules in terms of their tree characteristics and were therefore removed for the module conservation analysis. The separation between the actual and random modules is better and the phylogenetic signal is stronger in case of the dynamic pruning (see B and C for quantification). **F**) Total tree length and within-species diversity of the pruned and corresponding random modules for the dynamic and top50 pruning approaches. The separation between the actual and random modules is better and the contribution of the within-species diversity (confounding factors) to the total tree length is lower in case of the dynamic pruning (see D for quantification). **G-H**) Network conservation of HOXA2 based on dynamic and top50 pruning. In both cases, a linear model was fitted between the total tree lengths and within-species diversities across all modules, and the 95% prediction interval of the regression line (shaded in grey) was calculated. Modules that fell above the upper bound or below the lower bound of the prediction interval were considered diverged and conserved, respectively. Conservation of the HOXA2 module was captured using the dynamic, but not using the top50 approach. **B-D**) Asterisks indicate significance based on Wilcoxon tests: * *p <* 0.05, ** *p <* 0.01, *** *p <* 0.001.

**Supplementary Figure S5.**
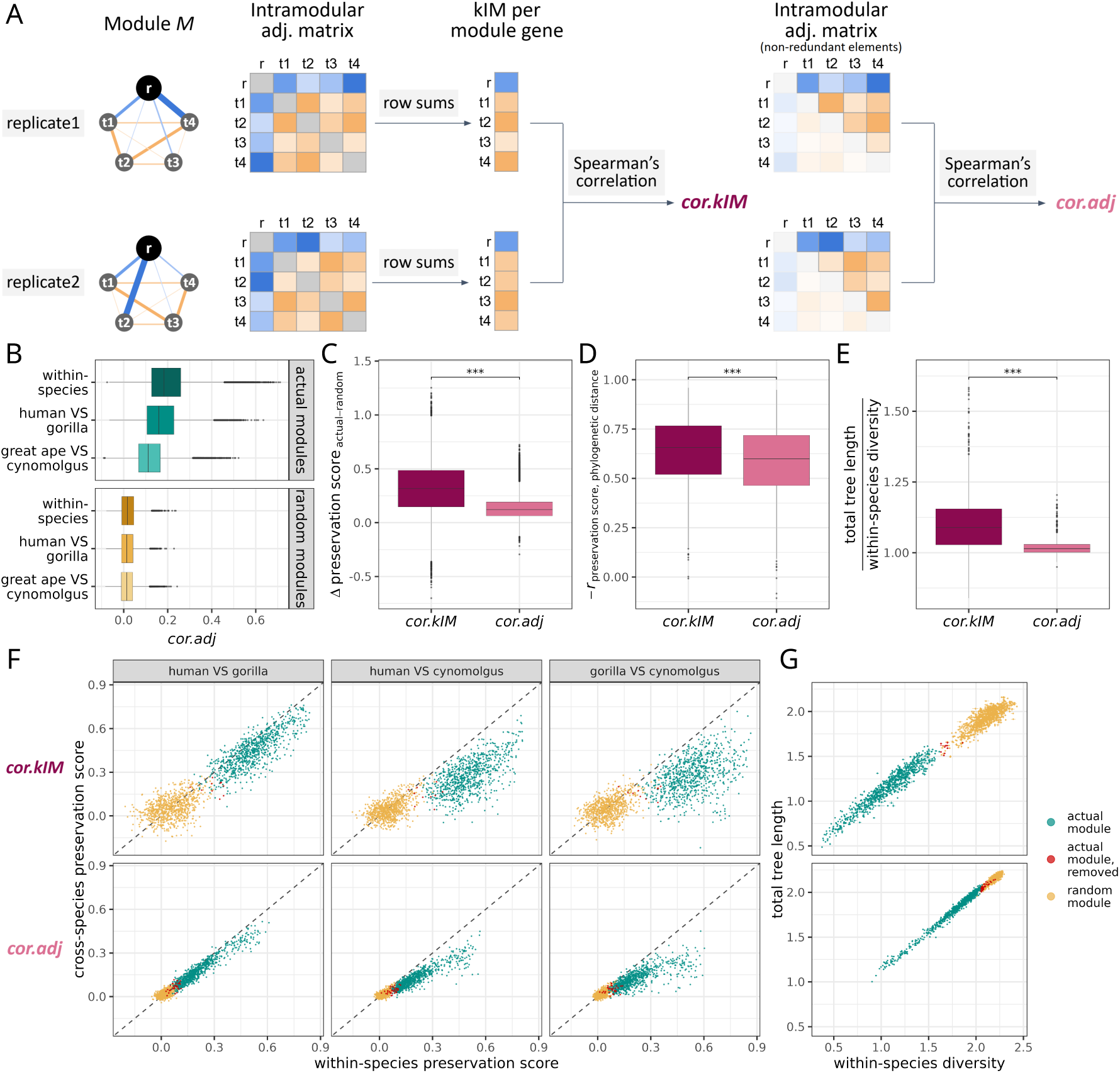
Comparison of the preservation statistics *cor.kIM* and *cor.adj*. **A**) Calculation of *cor.kIM* and *cor.adj*. **B**) Distribution of *cor.adj* for the pruned and corresponding random modules, split by divergence time of the replicates compared. **C**) Difference in the *cor.kIM* /*cor.adj* scores between each actual and the corresponding random module. **D**) Inverted Pearson’s correlation (−*r*) between *cor.kIM* /*cor.adj* and phylogenetic distance. **E**) Ratios of the total tree lengths and within-species diversities in the tree representations of the pruned modules, reconstructed based on the *cor.kIM* and *cor.adj* scores. **F**) Within-species and cross-species *cor.kIM* and *cor.adj* scores of the pruned and corresponding random modules, split by species pair. Actual modules in red were not distinct enough from the random modules in terms of their tree characteristics and were therefore removed for the module conservation analysis. The separation between the actual and random modules is better and the phylogenetic signal is stronger in case of *cor.kIM* (see C and D for quantification). **G**) Total tree length and within-species diversity of the pruned and corresponding random modules, based on the trees reconstructed using the *cor.kIM* and *cor.adj* scores. The separation between the actual and random modules is better and the contribution of the within-species diversity (confounding factors) to the total tree length is lower in case of *cor.kIM* (see E for quantification). **C-E**) Asterisks indicate significance based on Wilcoxon tests: * *p <* 0.05, ** *p <* 0.01, *** *p <* 0.001.

**Supplementary Figure S6.**
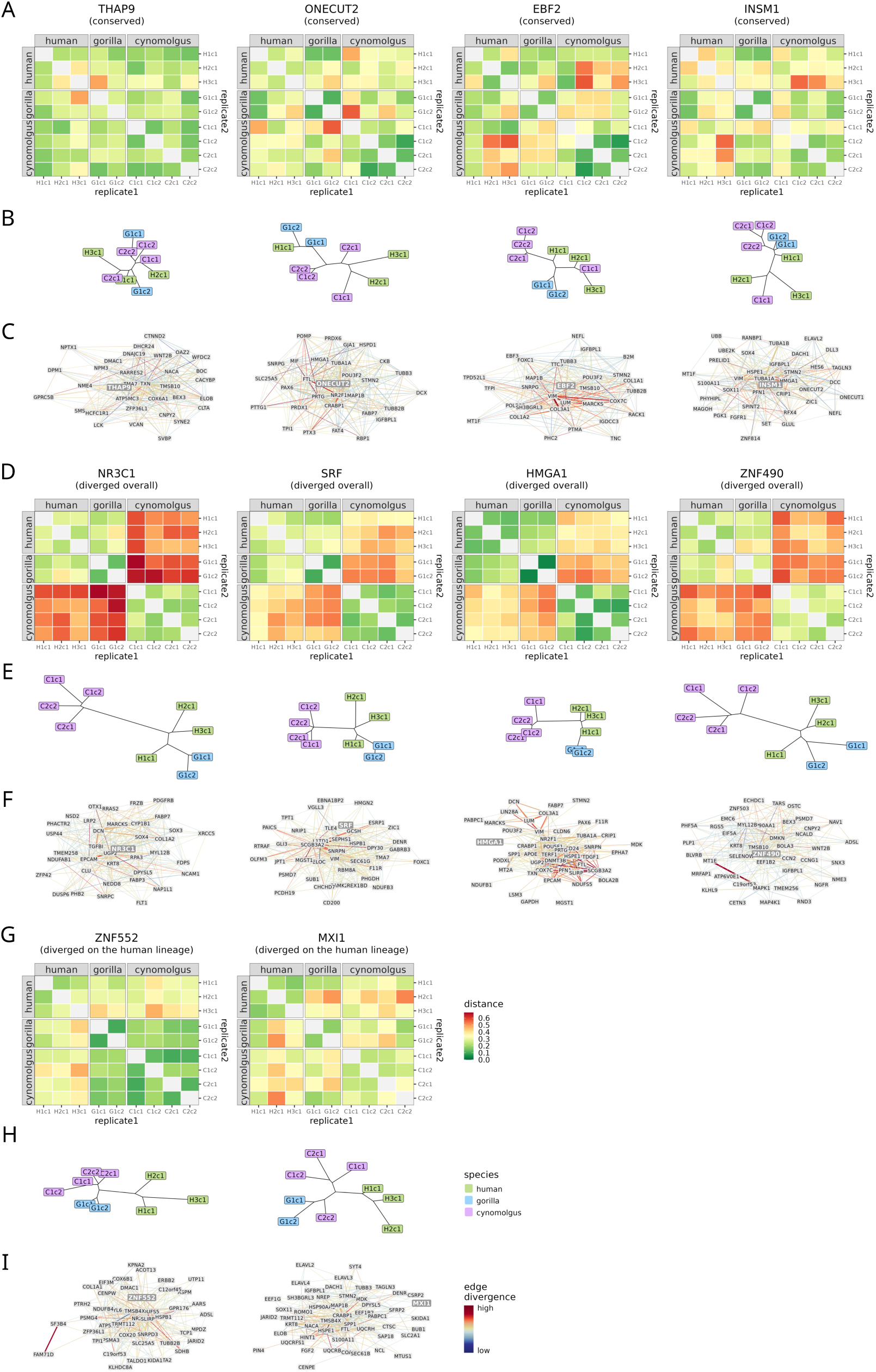
Distance matrices, tree reconstructions and network representations of the most conserved and most diverged modules. **A**) **D**) **G**) Distance matrices based on *cor.kIM* for the five most conserved modules, the five most diverged modules overall and the three diverged modules on the human lineage, respectively (the three example modules in Fig. 3F-H are excluded). **B**) **E**) **H**) Neighbor-joining trees for the same modules as in A, D, and G, respectively. **C**) **F**) **I**) The 200 strongest connections of the same modules as in A, D, and G, respectively. The edge thickness represents the consensus edge weights (scaled per module) and edge color represents how different the mean edge weights are across the three species (−log_10_*F* based on an ANOVA per edge).

**Supplementary Figure S7.**
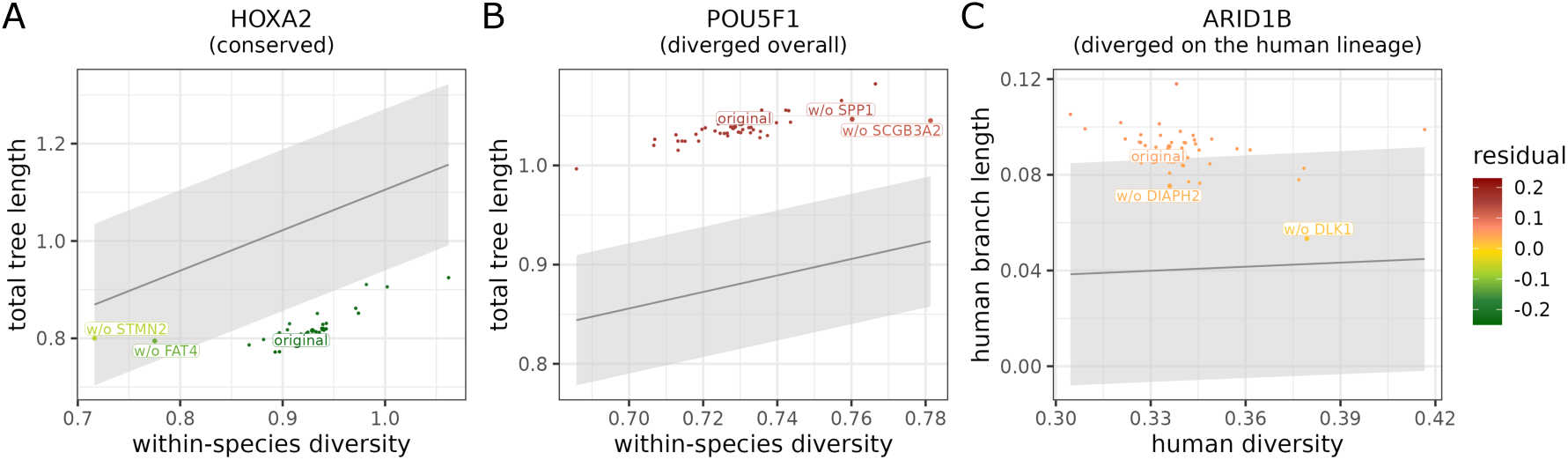
Identifying the top-contributing genes within the HOXA2, POU5F1 and ARID1B modules using jackknifing. **A-C**) In case of HOXA2 and POU5F1, the black line represents the regression line between the total tree lengths and within-species diversities of all modules, while in case of ARID1B, the black line represents the regression line between the human subtree lengths and human diversities of all human-monophyletic modules. The grey area marks the 95% prediction interval of the regression line in all cases. Each data point represents the original or a jackknifed version of the module, colored by its deviation from the regression line. The two jackknife versions with the highest reduction in the signal of conservation or divergence are labeled for each module. The labels indicate the target genes that have been removed. These are the top-contributing targets, i.e. in case of the conserved HOXA2 module, they contribute the most to the signal of conservation, while in case of the diverged POU5F1 and ARID1B modules, they contribute the most to the signal of divergence.

**Supplementary Figure S8.**
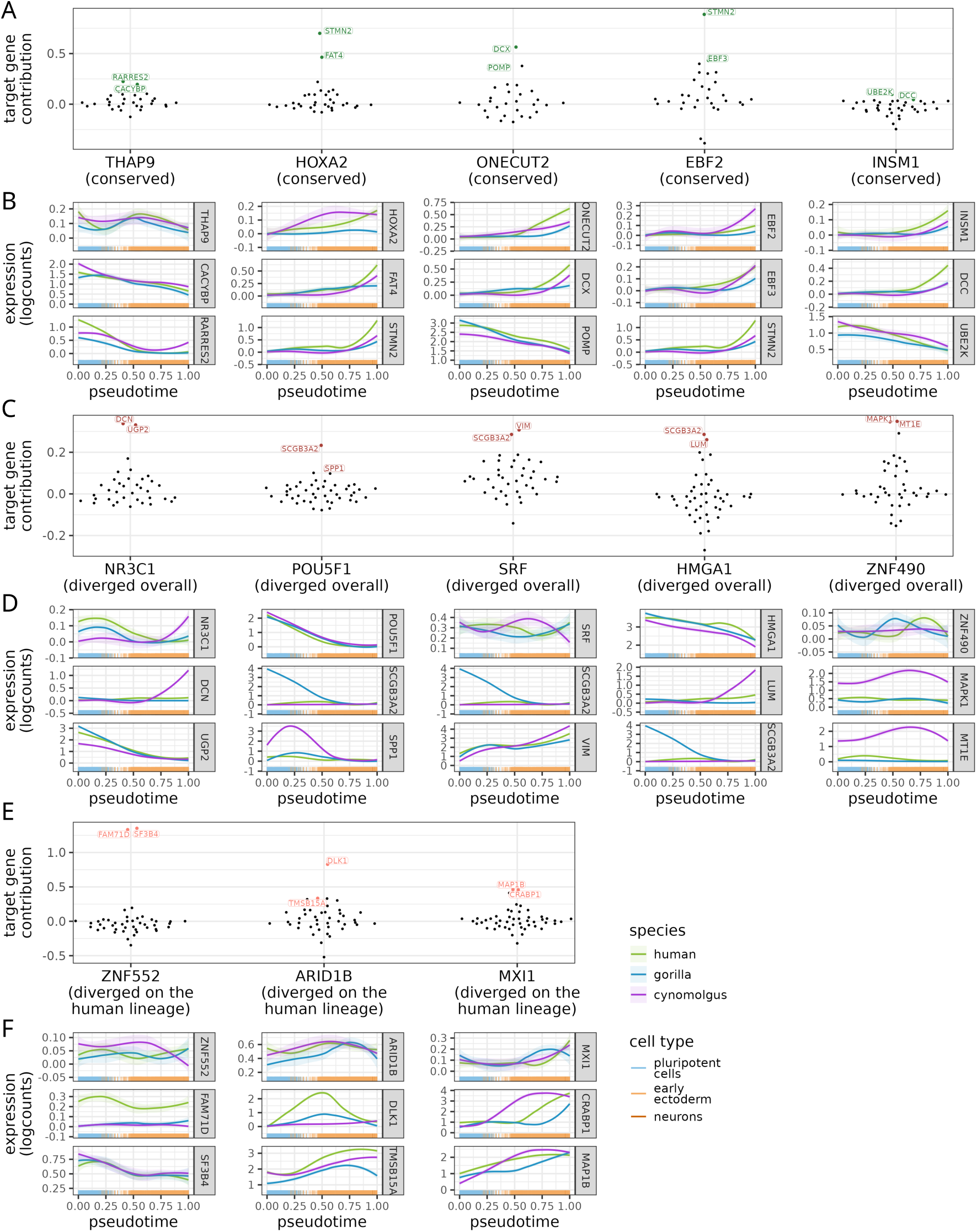
Target gene contributions and expression profiles for the most conserved and most diverged modules. **A**) **C**) **E**) Target gene contributions for the five most diverged modules overall, the five most conserved modules, and the three diverged modules on the human lineage, respectively. For conserved modules, top-contributing targets drive the conservation signal, whereas for diverged modules, they drive the divergence signal. **B**) **D**) **F**) Expression profiles of the central regulator and its two targets with the highest contribution scores for the five most diverged modules overall, the five most conserved modules, and the three diverged modules on the human lineage, respectively.

**Supplementary Figure S9.**
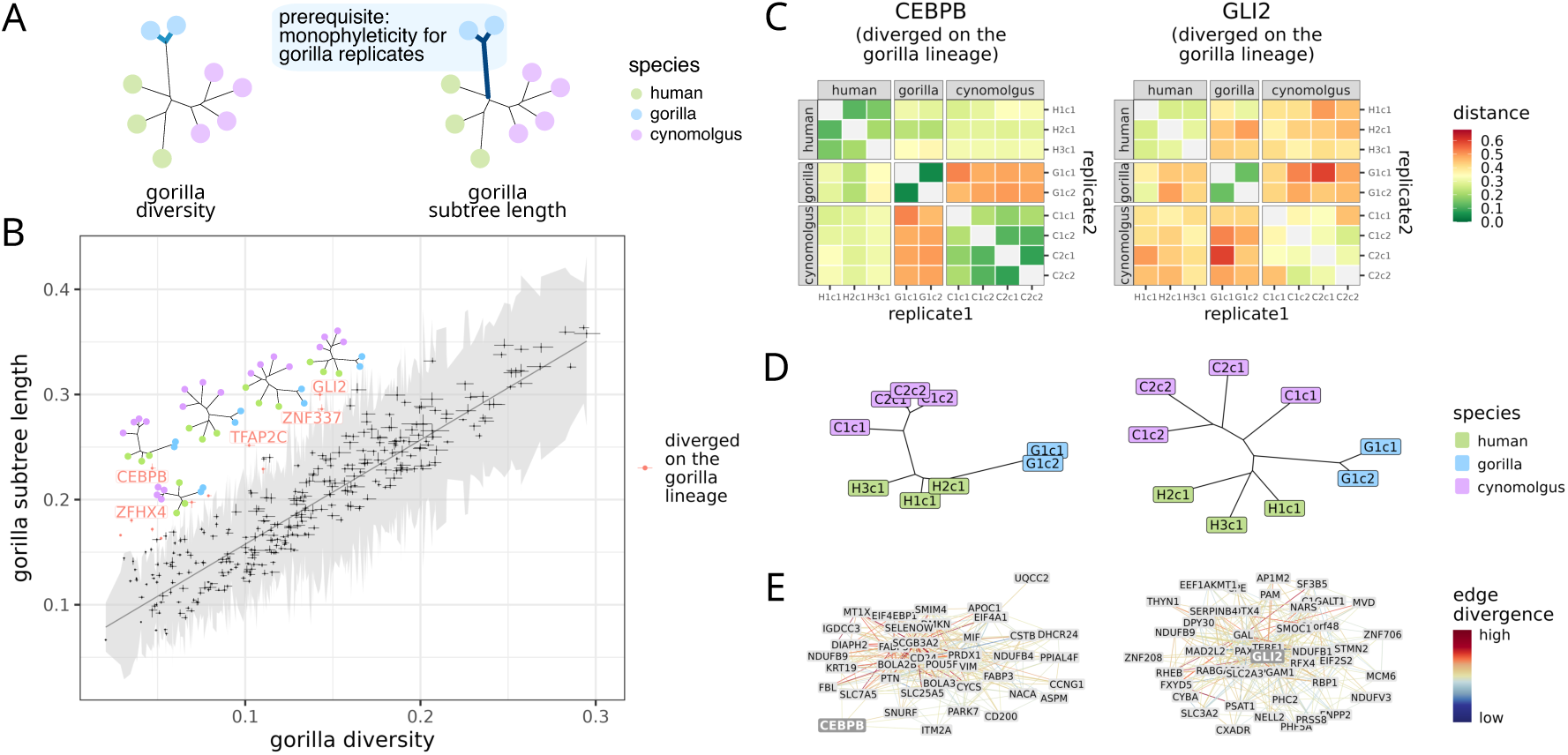
Divergence on the gorilla and cynomolgus lineages. **A**) Gorilla diversity and gorilla subtree length. The gorilla subtree length is only defined if the tree is robustly monophyletic for the gorilla replicates. **B**) Quantification of module divergence on the gorilla lineage. A linear model was fitted between the gorilla subtree lengths and gorilla diversities across all gorilla-monophyletic modules, and the 95% prediction interval of the regression line (shaded in grey) was calculated. Modules that fell above the upper bound of the prediction interval were considered diverged. The five most diverged modules are labeled and depicted using the tree representations. **C**) Distance matrices based on *cor.kIM* for the two most diverged modules on the gorilla lineage. **D**) Neighbor-joining trees for the two most diverged modules on the gorilla lineage. **E**) The 200 strongest connections of the two most diverged modules on the gorilla lineage. The edge thickness represents the consensus edge weights (scaled per module) and edge color represents how different the mean edge weights are across the three species (−log_10_*F* based on an ANOVA per edge).

**Supplementary Figure S10.**
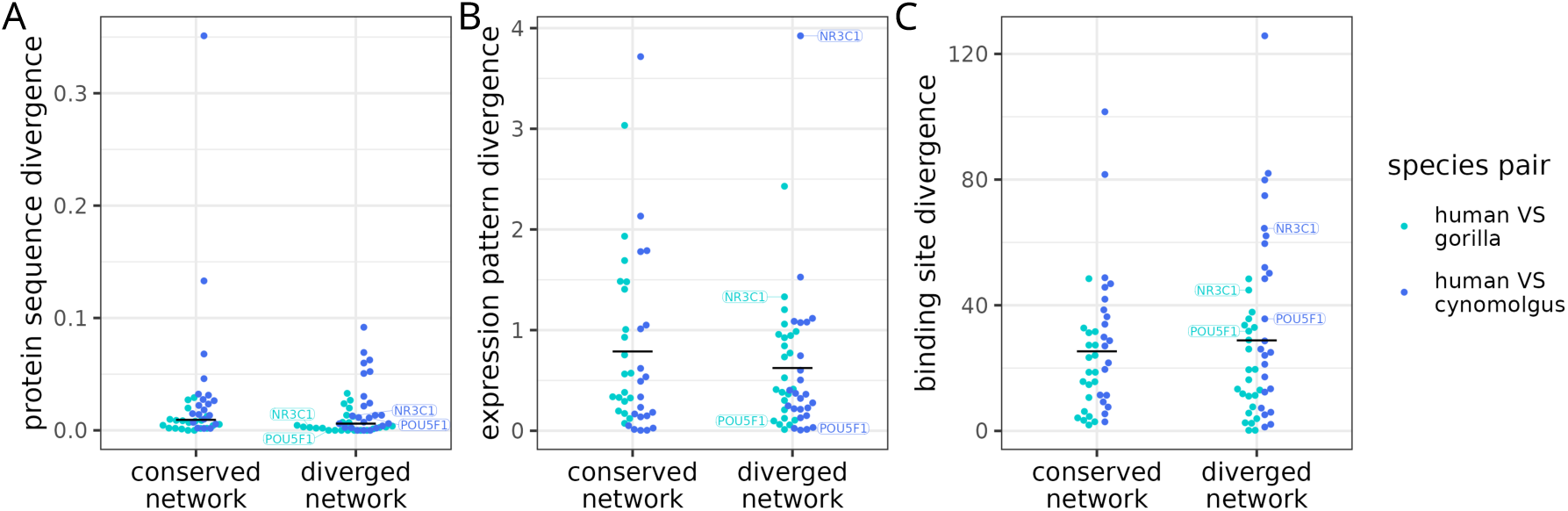
Relationship between module divergence and the sequence, expression pattern or binding site divergence of the master regulator. **A**) Sequence divergence of the regulators associated with conserved and diverged modules. The sequence divergence was quantified as the fraction of amino acid substitutions in the human-gorilla and human-cynomolgus protein sequence alignments. No significant difference was detected in this metric between regulators with conserved and diverged networks (tested using the linear model *sequence divergence* ∼ *network conservation category + phylogenetic distance*, *β* = 0.01, *p* = 0.24). **B**) Expression pattern divergence of the regulators associated with conserved and diverged modules. Expression pattern divergence was defined based on a DE analysis: log_2_ fold changes between the late and early pseudotime stages were estimated per species, then the absolute differences between the log_2_ fold change values, normalized by mean expression, were calculated for each species pair. No significant difference was detected in this metric between regulators with conserved and diverged networks (tested using the linear model *expression pattern divergence* ∼ *network conservation category + phylogenetic distance* for non-redundant species comparisons, *β* = −0.16, *p* = 0.33). **C**) Binding site divergence of the regulators associated with conserved and diverged modules. The binding site divergence was calculated based on scoring all annotated motifs of the regulators in the ATAC-seq peaks associated with their module member genes. For each regulator, the scores of non-redundant motifs were summed per gene, then the absolute differences of these sum motif scores were calculated per gene and species pair. Finally, these differences were summarized per species pair and module. No significant difference was detected in this metric between all regulators with conserved and diverged networks (tested using the linear model *binding site divergence* ∼ *network conservation category + phylogenetic distance* for non-redundant species comparisons, *β* = 3.48, *p* = 0.47). However, the difference was significant between the five most conserved and five most diverged modules (Fig. 3P). **A-C**) POU5F1 and NR3C1, the two main diverged examples, are labeled.

### Supplementary tables

**Supplementary Table 1. Experiments and cell lines.** Cell lines (replicates) used in the neural differentiation scRNA-seq, iPSC and NPC bulk ATAC-seq, iPSC and NPC long-read RNA-seq and CRISPRi experiments along with information about the species and individuals.

**Supplementary Table 2. Replicates in the primate brain dataset.** IDs of the donors (replicates) in the original Jorstad *et al.* paper and the shortened IDs used in this paper along with information about the species.

**Supplementary Table 3. Pruned modules in the neural differentiation dataset.** Regulators, module sizes and member genes of the 836 pruned modules assigned in the neural differentiation dataset, including the regulator-target adjacency and intramodular connectivity of each member gene, the replicates supporting each regulator-target edge, and the direction of correlation between each regulator-target pair.

**Supplementary Table 4. Pruned modules in the primate brain dataset.** Regulators, module sizes and member genes of the 499 pruned modules assigned in the primate brain dataset, including the regulator-target adjacency and intramodular connectivity of each member gene, and the direction of correlation between each regulator-target pair.

**Supplementary Table 5. Properties of the modules in the neural differentiation dataset.** Sequence-, expression-, binding site- and network-related characteristics of the 836 regulators as well as information about the overall and lineage-specific divergence of their network modules.

**Supplementary Table 6. Properties of the modules in the primate brain dataset.** Mean expression and network characteristics of the 499 regulators as well as information about the overall and lineage-specific divergence of their network modules.

**Supplementary Table 7. Target gene contribution scores in the neural differentiation dataset.** Contribution score of each target gene in the conserved and diverged modules of the neural differentiation dataset, and the underlying tree characteristics and residuals of the original and jackknifed versions of the modules.

**Supplementary Table 8. POU5F1-targeting gRNAs in the CRISPRi experiment.** IDs, sequences, cell numbers and knockdown efficiencies of the gRNAs designed to target POU5F1 in human and cynomolgus iPS cells. The best cross-species gRNA pair selected for the DE analysis is marked in the column ”best gRNA pair”.

**Supplementary Table 9. Control gRNAs in the CRISPRi experiment.** IDs, sequences and cell numbers of the non-targeting control gRNAs designed for human and cynomolgus iPS cells.

**Supplementary Table 10. POU5F1 perturbation outcome in human and cynomolgus iPS cells.** Results of the DE analysis contrasting perturbed and control human cells (DE human), perturbed and control cynomolgus cells (DE cynomolgus) and the perturbed-control differences across the two species (DR).

### Supplementary methods

#### Neural differentiation experiment

Primate iPSCs [66, 67, 69, 70] (Supplementary Table 1) were cultured in StemFit + bFGF [67] and differentiated into NPCs via dual-SMAD inhibition [71, 72] over the course of 9 days. On days 0, 1, 3, 5, 7 and 9, the progression of the differentiation was validated by OCT4 and PAX6 staining, and cells were sampled for scRNA-seq. Libraries were prepared using the mcSCRB-seq protocol [73] and sequenced on an Illumina HiSeq 1500 instrument with 2×100 bp paired-end reads.

#### Processing of the differentiation dataset

After polyA trimming, the FASTQ data were processed using the zUMIs pipeline [74] that utilizes the aligner STAR [94]. We mapped all reads to three reference genomes: GRCh38/hg38 (GENCODE release 32), Kamilah GGO v0/gorGor6 (UCSC, Aug. 2019) and Macaca fascicularis 6.0/macFas6 (ENSEMBL release 109). To reduce computational time, we removed contigs smaller than 150 kb from the gorGor6 genome. The gorilla and macaque GTF files were created by Liftoff of the hg38 annotation to the corresponding primate genomes [25], followed by removal of transcripts with partial mapping (*<*50%), low sequence identity (*<*50%) or excessive length (*>*100 bp difference and *>*2 length ratio).

We assigned the cells to the three species based on their barcode information and set filters for cell quality based on 1) the number of UMIs (2,000 *<* # of UMIs *<* 30,000 for human, 1,000 *<* # of UMIs *<* 32,000 for gorilla, and 1,000 *<* # of UMIs *<* 34,000 for cynomolgus), 2) the number of genes (*>*700), 3) the percentage of mitochondrial reads (*<*6%), and 4) the percentage of spike-in UMIs (*<*20%). We kept genes present in all three annotation files and expressed in *>*1% of the cells in all replicates of at least one species. In addition, we removed all mitochondrial, non-protein coding, ribosomal, histone and pseudogenes. Histone, ribosomal and pseudogenes were defined using the gene group and locus group annotations from the HGNC database [95]. We summed up the counts for paralog groups with *>*95% mean sequence identity based on the *Hsapiens gene ensembl* dataset of BioMart [96] and labelled each group by the first gene name among the paralogs in alphabetical order.

We normalized the count matrices using scran::computeSumFactors [75] and scran:: computeSumFactors [97]. For certain lines of analysis, we additionally applied per-batch scaling normalization using batchelor::multiBatchNorm [78]. We assigned cell type labels via SingleR [76] using a human embryoid body dataset [77] as reference. Non-neuronal cell types (Neural Crest, Mesoderm, Endothelial Cells, Endoderm), as well as the day7 cells from the cynomolgus replicates C1c1 and C1c2 were removed for further analysis. For pseudotime inference, we regressed out the batch effects due to replicate differences using batchelor::fastMNN [78], then inferred a pseudotime trajectory using SCORPIUS [79] on a low-dimensional embedding of the batch-corrected counts (dist = ’spearman’ and ndim = 25).

#### Network inference for the differentiation dataset

For the network inference, genes expressed in only 1 or 2 cells of a replicate were set to zero in those few cells as well to avoid spurious connections in the network. Raw counts were then normalized per replicate via transformGamPoi::transformGamPoi, using the randomized quantile residual transformation.

Based on the normalized counts, networks were reconstructed per replicate with the help of GRNBoost2. As the list of transcriptional regulators, we used all 11,630 genes in the filtered count matrix. We ran GRNBoost2 [16] using the multiprocessing implementation of Arboreto (arboreto with multiprocessing.py) included in pySCENIC [81], 10 times for each replicate with 10 different seeds to make the results less dependent on the stochastic nature of the algorithm. We loaded and processed the networks using igraph [98].

First, we summarized the results per replicate. For each replicate, a given edge could appear up to 20 times (10 runs × 2 directions) in the output. For edges that appeared once or not at all, we set the edge weight to 0. For edges that appeared at least twice, we defined the edge weight as the mean importance score across the 20 occurrences; in case an edge was not present for one of these 20 predictions, this was factored into the mean as a 0. The edge weights were then rescaled between 0 and 1 for all replicates together. To avoid false positive edges due to mapping errors, we set the weight of an edge to 0, if it connected two genes that had overlapping annotations in any of the species’ genomes. The adjacencies resulting from this network inference and processing approach were undirected (a single value per gene pair; regulator and target are not distinguished) and unsigned (strong activating and repressing interactions are both characterized by high adjacencies).

#### Selection of the central regulators

We modeled the mean-variance relationship of the log-normalized counts and decomposed the variance into technical and biological components per species using the function scran::modelGeneVar [75], then identified the genes that had a positive biological variance in at least one species. We intersected these genes with transcriptional regulators that have at least one known motif based on the JASPAR 2024 core [18], JASPAR 2024 unvalidated [18] and the IMAGE databases [19].

#### Module-level summaries of the pruning metrics

The replicate-wise networks were aggregated into a consensus network, and based on the consensus adjacencies, modules were assigned around each central regulator as described in the Methods.

The pruning metrics can be summarized per module as the mean regulator-target adjacency and the mean size-corrected kIM (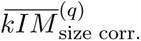, Supplementary Table 5):

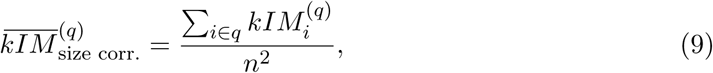

where *n* is the module size, i.e. the total number of genes in the module.

#### Module overlaps

In the neural differentiation dataset, the overlap between the member genes of two modules *q*_1_ and *q*_2_ was quantified using the overlap fraction:

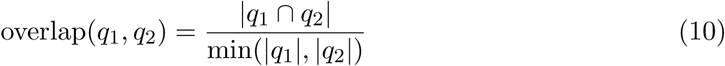

Protein-protein interactions between the regulators were identified using STRINGdb’s human database [28] with a minimum score threshold of 200. We found that interacting regulators had significantly higher target gene overlaps between their pruned modules than non-interacting regulators (Wilcoxon test, *n*_1_ = 17, 670, *n*_2_ = 331, 360, *p <* 1 · 10^−16^, Supplementary Fig. S2B). This was also the case for the initial modules, although to a lesser extent, and could not be observed at all for the random modules.

#### Pathway enrichment analysis

For the neural differentiation dataset, we performed pathway enrichment analysis on the activated targets of the initial, pruned and random modules. To quantify enrichment, we used Reactome [42, 43], with all genes in the network as universe. We regarded pathways with an adjusted *p*-value *<*0.1 (Benjamini-Hochberg FDR correction) as significant.

#### Module eigengenes

To assign a directionality to each non-zero edge in the consensus networks, we correlated the expression profiles of the corresponding gene pairs. We calculated a modified Spearman’s rho using scran::correlatePairs [75] with replicates as blocks.

Next, we summarized the expression profiles of the modules using the concept of module eigengenes. We selected the module genes that showed a positive correlation to the central regulator, subsetted the normalized count matrix for these genes, then calculated the 1st principal component of these matrices aligned with the average expression as implemented by WGCNA::moduleEigengenes[41].

#### ATAC-seq data

Part of the ATAC-seq dataset used for motif enrichment analysis has been previously published [69]; here we describe both the published and unpublished work. ATAC-seq data were collected from seven primate cell lines [67–70] (Supplementary Table 1): two human iPSC clones of two individuals (H1c2, H2c1), two gorilla iPSC clones of one individual (G1c2, G1c3) and three cynomolgus iPSC clones of two individuals (C1c1, C1c3, C2c1). Two human clones (H1c2, H2c1) and two cynomolgus clones (C1c1, C2c1) have been differentiated as described in section Neural differentiation experiment and ATAC-seq data were also collected from NPCs after differentiation. All libraries were generated using the Omni-ATAC protocol [86] with minor modifications.

We mapped reads to the same reference genomes as for the scRNA-seq data using BWA-MEM2 [87]. We called peaks jointly for all replicates per cell type and species using genrich [88] in a two-step process. In the first step, a blacklist was created for each genome by taking peaks with unusually high widths and signal values. In the second step, peak calling was repeated using these blacklists. To gain information about accessibility in gorilla NPCs, the orthologous regions for the human NPC peaks were identified in the gorGor6 genome using the UCSC LiftOver tool [89].

#### Long-read RNA-seq data

Total RNA was isolated from samples of seven primate cell lines [67–69] (Supplementary Table 1): two human iPSC samples (H2c1, H2c2), a human NPC sample (H1c2), a gorilla iPSC sample (G1c1), a gorilla NPC sample (G1c2), a cynomolgus iPSC sample (C1c1), and three cynomolgus NPC samples (two different passages of C1c1, and C1c2). For all cell lines, 50 ng RNA were reverse-transcribed using the PCR-cDNA Barcoding Kit SQK-PCB109 (Oxford Nanopore Technologies). Purified barcoded cDNAs were pooled and sequenced on a PromethION R9.4.1 flowcell (Oxford Nanopore Technologies).

Reads were mapped to the same reference genomes as for the scRNA-seq data using minimap2 [99]. Aligned reads were converted into transcript models using the pinfish pipeline (Oxford Nanopore Technologies). Gene names were assigned to the transcripts if *>*60% of their base pairs across all exons overlapped with exons of the respective gene in the GENCODE or liftoff annotation.

#### Peak-to-gene associations

We identified the transcriptional start sites (TSS) of active genes in human, gorilla and cynomolgus iPSCs and NPCs. TSS were defined as the 5’-most positions of transcript models derived from the GENCODE or liftoff annotations, and from long-read RNA-seq data of the species and cell type of interest. TSS within 100 bp of each other were subsequently merged. A gene was regarded as active if it fulfilled at least one of the following criteria: 1) the TSS was identified based on long read RNA-seq data, 2) the TSS overlapped with an ATAC-seq peak of the corresponding species and cell type, 3) the gene was expressed in *>*5% of the cells and had a mean log-expression of *>*0.05 in the scRNA-seq data in the corresponding species and cell type.

For each cell type and species, ATAC-seq peaks were associated with genes based on their distance to TSSs as described in [69]. Peaks within 2 kb of a TSS were regarded as promoters and were associated with all TSSs within that distance. All other peaks within 20 kb of a TSS were considered enhancers and were associated with the two closest TSSs in each direction, unless the distance to one TSS was *>*10× smaller than to the other TSS; in that case, only the closest TSS was kept.

Next, peak-to-gene associations from iPSCs and NPCs of the same species were unified. If an iPSC and NPC peak overlapped by more than 50%, and the associated genes (in either direction) differed between cell types, the gene closest to the peak was assigned in both cases.

#### Motif scoring

For each of the 836 regulators in the neural differentiation dataset, we identified known binding motifs and their position weight matrices from the JASPAR 2022 vertebrate core [90], JASPAR 2022 vertebrate unvalidated [90] and IMAGE databases [19]. We scored all motifs of a regulator in all peaks associated with the genes in its initial, pruned and random modules using Cluster-Buster [91]. We used a padding of 500 bp and a range of 10,000 bp to calculate local nucleotide abundances and kept all hits without a motif score threshold.

Next, we collapsed redundant motif hits. If several hits on the same strand, within the same peak and for the same regulator overlapped, we kept only the one with the highest score. If an iPSC and NPC peak of the same species overlapped, we merged them and performed this summarization across the entire union peak. Finally, we summarized the binding potential per gene, regulator and species (*G*) by summing up the non-redundant motif scores (*S*) across all associated peaks in the given species:

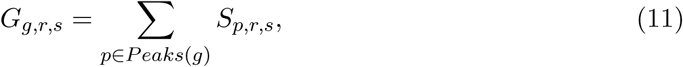

where *g* is the gene, *r* is the regulator, *s* is the species, and *Peaks*(*g*) is the set of peaks associated to *g*. For genes that had associated peaks but no motif hits, this summarized score was regarded as 0, while genes that had no associated peaks to begin with were excluded from the downstream analysis.

#### Module validation using motif scores

To compare the random, initial and pruned modules of the neural differentiation dataset in terms of binding potential of the regulator, we further summarized the scores per module and species (*M*) by taking the median across all module member genes (Supplementary Table 5):

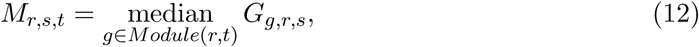

where *t* is the module type (initial, pruned or random), and *Module*(*r, t*) is the set of genes in the module of type *t* assigned to regulator *r*.

Then we also calculated the Δ motif scores between the corresponding initial and random as well as between the corresponding pruned and random modules. Finally, we performed paired two-sided Wilcoxon tests per species to compare 1) the initial and random scores (*M_r,s,_*_initial_ vs. *M_r,s,_*_random_), 2) the pruned and random scores (*M_r,s,_*_pruned_ VS *M_r,s,_*_random_), and 3) the pruned-random and initial-random Δ scores (*M_r,s,_*_pruned_ − *M_r,s,_*_random_ VS *M_r,s,_*_initial_ − *^M^_r,s,_*_random_^).^

#### Cross-species differences in motif scores

To investigate cross-species differences in binding potential, we calculated the differences of gene-wise motif scores between pairs of species (Δ*G*, Supplementary Table 5):

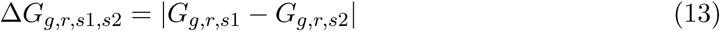

Then we summarized the Δ*G* values per pruned module and species pair (Δ*M*) by taking the median across all genes to obtain a measure of binding site divergence between the two species:

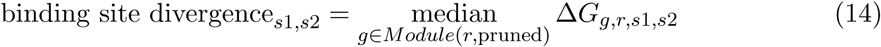

Finally, to test whether regulators with conserved and diverged network modules differ in terms of their binding site divergence, we subsetted non-redundant species comparisons (human-gorilla and human-cynomolgus only) and fitted the following linear model across the regulators with significant conservation or divergence detected in the CroCoNet analysis:

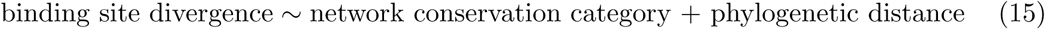

Based on the model, regulators of diverged modules tend to be associated with higher binding site divergence than regulators of conserved modules, but this difference is not significant (*β* = 3.48, *p* = 0.47). When comparing the five most diverged and five most conserved modules only, this difference is more pronounced, reaching statistical significance (*β* = 14.9, *p* = 0.024).

#### POU5F1 ChIP-seq data

To validate the POU5F1 module assignment in the neural differentiation dataset, we used POU5F1 ChIP-seq data from human primed embryonic stem cells [27]. We mapped the reads from both output replicates and the input control to the hg38 reference genome using bowtie [100]. We called peaks jointly for the two replicates using genrich [88] with a blacklist provided by ENCODE [101].

For each gene in the network, we checked whether any of the associated ATAC-peaks overlapped with any of the POU5F1 ChIP-seq peaks. We found that POU5F1 module member genes showed such an overlap significantly more often than all other genes (two-sided Fisher’s exact test, *p <* 0.001).

#### Preservation statistics

To introduce the two preservation statistics that CroCoNet relies on, let *q* denote a consensus module with *n_q_* genes, and let *A*^[*net*](*q*)^ denote its intramodular adjacency matrix, i.e. the *n_q_* × *n_q_* dimensional subset of the entire adjacency matrix in the network *net* that contains only the module genes. In addition, let *vectorizeMatrix()* denote a function that takes a matrix as input and returns its non-redundant elements (i.e. the upper triangular elements in case of a symmetric matrix) in a vector form.

Then the correlation of adjacencies (*cor.adj*) and correlation of intramodular connectivities (*cor.kIM*) [7] can be calculated between the networks of two replicates (*net1* and *net2*) as follows:

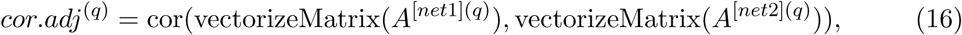

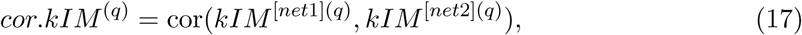

#### Edge divergence

For each edge in the network, we calculated an edge divergence score based on the edge weights inferred in the different species and replicates. We performed a one-way ANOVA with the species as groups and the replicates as observations, calculated the corresponding *F* -statistic and defined the edge divergence as follows:

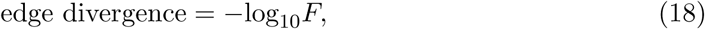

#### Processing of the POU5F1 CRISPRi data

Reads were processed using 10x CellRanger (Version 7.0.0, https://support.10xgenomics.com/single-cell-gene-expression/software/pipelines/latest/what-is-cell-ranger). mapped reads of the GEX library to the hg38 and macFas6 reference genomes extended by the sequence of the dCas9-construct, and reads of the CRISPR library to a custom reference created using the gRNA protospacer sequences. The reads were demultiplexed into species and individuals in a two-step process using cellsnp-lite [102] and vireo [103], based on a list of single nucleotide polymorphisms (SNPs) compiled from bulk RNA-seq data of the wild-type cell lines.

We kept cells that passed basic quality control, carried a single dominant gRNA (i.e. all other detected gRNAs in the cell made up *<*10% of gRNA UMIs and *<*1000 UMIs combined), and had *>*10 UMIs supporting the dominant gRNA. We excluded gRNAs from a species if they were detected in *<*20 cells in either individual of that species. To remove control gRNAs with an unwanted transcriptomic effect, we performed differential expression (DE) analysis by limma-trend [104], comparing each control gRNA against all others. We iteratively removed any gRNA that had more DE genes (adjusted *p*-value *<*0.05) than *mean* + 3*σ* across the DE gene counts of all control gRNAs. We repeated this process until all remaining gRNAs fell within this range. After these filtering steps, we retained 1,635 POU5F1-perturbed cells (5 gRNAs) and 16,981 control cells (39 gRNAs) for the human, and 1,482 POU5F1-perturbed cells (6 gRNAs) and 16,421 control cells (37 gRNAs) for the cynomolgus (Supplementary Tables 8-9).

We kept genes that could be transferred from the hg38 to the macFas6 genome via Liftoff [25], and were expressed in at least one condition (perturbed or control) of at least one species (*>*10% of cells and *>*10 cells) robustly across replicates. In addition, we excluded non-protein coding, mitochondrial and Y-chromosomal genes. This filtering approach resulted in 10,413 genes retained.

To characterize the pluripotency state of the cells, we calculated a transcriptome-based stemness index for each cell. We applied a one-class logistic regression model [105] trained on pluripotent stem cell samples [106, 107]. The resulting scores were scaled between 0 and 1 across both species.

For visualization, we performed normalization, feature selection and PCA reduction using Seurat v5 [108], then integrated the data across individuals and batches per species using Harmony [109] with *θ* = 3, and finally applied UMAP dimensionality reduction using Seurat v5.

To quantify how well the gRNAs downregulate POU5F1, we calculated a % knockdown (% KD) value for each gRNA:

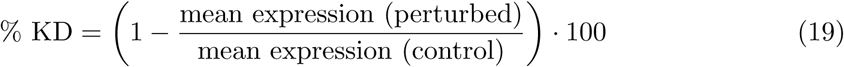

#### Protein sequence divergence

Hg38-gorGor6 and hg38-macFas6 protein alignments created by TOGA [48] were downloaded from http://genome.senckenberg.de/download/TOGA/human_hg38_reference/Primates/. For each of the 836 regulators in the neural differentiation dataset, we identified the longest canonical coding sequence (CCDS), or if no CCDS were annotated, the longest coding sequence (CDS) in the hg38 GENCODE v32 annotation. Based on the protein alignments corresponding to these CDS, we calculated a measure of protein sequence divergence as follows (Supplementary Table 5):

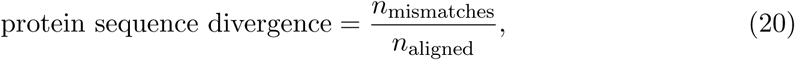

where *n*_aligned_ is the number of aligned amino acid positions where both species have a non-gap character, and *n*_mismatches_ is the number of mismatches among those aligned positions.

To test whether regulators with conserved and diverged network modules differ in terms of their protein sequence divergence, we fitted the following linear model across the regulators with significant conservation or divergence detected in the CroCoNet analysis:

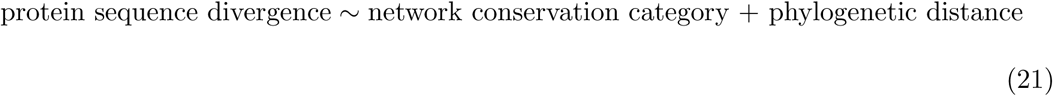

Based on the model, there is no significant difference in protein sequence divergence between regulators with conserved and diverged modules (*β* = 0.01, *p* = 0.24).

#### Expression pattern divergence

Based on pseudotime (*pt*), cells of the neural differentiation dataset were binned into three stages: early/iPSC (*pt* ≤ 0.25), intermediate (0.25 *< pt* ≤ 0.75), and late/NPC (*pt >* 0.75). The data were subsampled to an equal number of cells across the three species in each of the stages (early stage: 346 cells/species, intermediate stage: 155 cells/species, late stage: 223 cells/species). During the subsampling, we ensured that not only the number of cells but also the pseudotime distribution of each stage is as similar as possible across species.

Using dream [93], we fitted the following mixed effects model to the log-normalized counts of each gene:

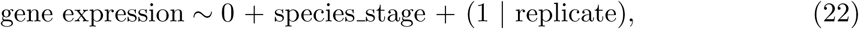

where ”species stage” denotes the combination of species and stage, which can take up 9 different values (3 species × 3 stages).

Based on the model, we estimated log_2_ fold changes between the late and early stages in each species. Subsequently, we calculated the absolute differences between the log_2_ fold change values per species pair (corresponding to estimates of a species:stage interaction term) normalized by the mean expression to obtain a measure of expression pattern divergence (Supplementary Table 5):

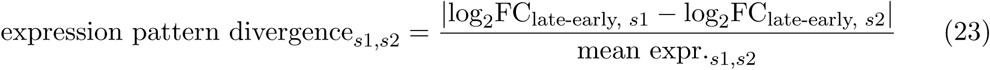

To test whether regulators with conserved and diverged network modules differ in terms of their expression pattern divergence, we subsetted non-redundant species comparisons (human-gorilla and human-cynomolgus only) and fitted the following linear model across the regulators with significant conservation or divergence detected in the CroCoNet analysis:

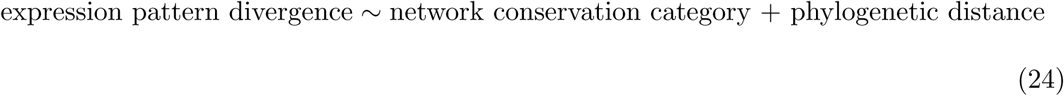

Based on the model, there is no significant difference in expression pattern divergence between regulators with conserved and diverged modules (*β* = −0.16, *p* = 0.33).

## References

1. Habib, N., Wapinski, I., Margalit, H., et al. A functional selection model explains evolutionary robustness despite plasticity in regulatory networks. Mol. Syst. Biol. 8, 619 (2012).

2. Krieger, G., Lupo, O., Levy, A. A., et al. Independent evolution of transcript abundance and gene regulatory dynamics. Genome Res. 30, 1000–1011 (2020).

3. Ovens, K., Eames, B. F. & McQuillan, I. Comparative Analyses of Gene Co-expression Networks: Implementations and Applications in the Study of Evolution. Front. Genet. 12, 695399 (2021).

4. Langfelder, P. & Horvath, S. Eigengene networks for studying the relationships between co-expression modules. BMC Syst. Biol. 1, 54 (2007).

5. Aibar, S., Gonźalez-Blas, C. B., Moerman, T., et al. SCENIC: single-cell regulatory network inference and clustering. Nat. Methods 14, 1083–1086 (2017).

6. Sousa, A. M. M., Zhu, Y., Raghanti, M. A., et al. Molecular and cellular reorganization of neural circuits in the human lineage. Science 358, 1027–1032 (2017).

7. Langfelder, P., Luo, R., Oldham, M. C., et al. Is my network module preserved and reproducible? PLoS Comput. Biol. 7, 1001057 (2011).

8. Gao, S., Wu, Z., Feng, X., et al. Comprehensive network modeling from single cell RNA sequencing of human and mouse reveals well conserved transcription regulation of hematopoiesis. BMC Genomics 21, 849 (2020).

9. True, J. R. & Haag, E. S. Developmental system drift and flexibility in evolutionary trajectories. Evol. Dev. 3, 109–119 (2001).

10. Fiers, M. W. E. J., Minnoye, L., Aibar, S., et al. Mapping gene regulatory networks from single-cell omics data. Brief. Funct. Genomics 17, 246–254 (2018).

11. Wills, Q. F., Livak, K. J., Tipping, A. J., et al. Single-cell gene expression analysis reveals genetic associations masked in whole-tissue experiments. Nat. Biotechnol. 31, 748–752 (2013).

12. Trapnell, C. Defining cell types and states with single-cell genomics. Genome Res. 25, 1491–1498 (2015).

13. Chen, S. & Mar, J. C. Evaluating methods of inferring gene regulatory networks highlights their lack of performance for single cell gene expression data. BMC Bioin-formatics 19, 1–21 (2018).

14. Pratapa, A., Jalihal, A. P., Law, J. N., et al. Benchmarking algorithms for gene regulatory network inference from single-cell transcriptomic data. Nat. Methods 17, 147–154 (2020).

15. McCalla, S. G., Fotuhi Siahpirani, A., Li, J., et al. Identifying strengths and weaknesses of methods for computational network inference from single-cell RNA-seq data. G3 13, jkad004 (2023).

16. Moerman, T., Santos, S. A., Gonźalez-Blas, C. B., et al. GRNBoost2 and Arboreto: efficient and scalable inference of gene regulatory networks. Bioinformatics 35, 2159–2161 (2019).

17. Kang, Y., Thieffry, D. & Cantini, L. Evaluating the Reproducibility of Single-Cell Gene Regulatory Network Inference Algorithms. Front. Genet. 12, 617282 (2021).

18. Rauluseviciute, I., Riudavets-Puig, R., Blanc-Mathieu, R., et al. JASPAR 2024: 20th anniversary of the open-access database of transcription factor binding profiles. Nucleic Acids Res 52, D174–D182 (2023).

19. Madsen, J. G. S., Rauch, A., Van Hauwaert, E. L., et al. Integrated analysis of motif activity and gene expression changes of transcription factors. Genome Res. 28, 243–255 (2018).

20. McDonald, J. H. & Kreitman, M. Adaptive protein evolution at the Adh locus in Drosophila. Nature 351, 652–654 (1991).

21. Hudson, R. R., Kreitman, M. & Aguadé, M. A test of neutral molecular evolution based on nucleotide data. Genetics 116, 153–159 (1987).

22. Jorstad, N. L., Song, J. H. T., Exposito-Alonso, D., et al. Comparative transcriptomics reveals human-specific cortical features. Science 382, eade9516 (2023).

23. Suresh, H., Crow, M., Jorstad, N., et al. Comparative single-cell transcriptomic analysis of primate brains highlights human-specific regulatory evolution. Nat Ecol Evol 7, 1930–1943 (2023).

24. Frankish, A., Diekhans, M., Jungreis, I., et al. GENCODE 2021. Nucleic Acids Res. 49, D916–D923 (2021).

25. Shumate, A. & Salzberg, S. L. Liftoff: accurate mapping of gene annotations. Bioin-formatics 37, 1639–1643 (2021).

26. Jocher, J., Janssen, P., Vieth, B., et al. Identification and comparison of orthologous cell types from primate embryoid bodies shows limits of marker gene transferability. eLife 14, RP105398 (2025).

27. Ji, X., Dadon, D. B., Powell, B. E., et al. 3D Chromosome Regulatory Landscape of Human Pluripotent Cells. Cell Stem Cell 18, 262–275 (2016).

28. Szklarczyk, D., Gable, A. L., Lyon, D., et al. STRING v11: protein–protein association networks with increased coverage, supporting functional discovery in genome-wide experimental datasets. Nucleic Acids Res. 47, D607–D613 (2019).

29. Chen, X., Xu, H., Yuan, P., et al. Integration of external signaling pathways with the core transcriptional network in embryonic stem cells. Cell 133, 1106–1117 (2008).

30. Van den Berg, D. L. C., Snoek, T., Mullin, N. P., et al. An Oct4-centered protein interaction network in embryonic stem cells. Cell Stem Cell 6, 369–381 (2010).

31. Boyer, L. A., Lee, T. I., Cole, M. F., et al. Core transcriptional regulatory circuitry in human embryonic stem cells. Cell 122, 947–956 (2005).

32. Chia, N.-Y., Chan, Y.-S., Feng, B., et al. A genome-wide RNAi screen reveals determinants of human embryonic stem cell identity. Nature 468, 316–320 (2010).

33. Ying, Q. L., Nichols, J., Chambers, I., et al. BMP induction of Id proteins suppresses differentiation and sustains embryonic stem cell self-renewal in collaboration with STAT3. Cell 115, 281–292 (2003).

34. Pasini, D., Cloos, P. A. C., Walfridsson, J., et al. JARID2 regulates binding of the Polycomb repressive complex 2 to target genes in ES cells. Nature 464, 306–310 (2010).

35. Pevny, L. & Placzek, M. SOX genes and neural progenitor identity. Curr. Opin. Neurobiol. 15, 7–13 (2005).

36. Kageyama, R., Ohtsuka, T., Shimojo, H., et al. Dynamic Notch signaling in neural progenitor cells and a revised view of lateral inhibition. Nat. Neurosci. 11, 1247–1251 (2008).

37. Lee, H.-K., Lee, H.-S. & Moody, S. A. Neural transcription factors: from embryos to neural stem cells. Mol. Cells 37, 705–712 (2014).

38. Huang, B., Li, X., Tu, X., et al. OTX1 regulates cell cycle progression of neural progenitors in the developing cerebral cortex. J. Biol. Chem. 293, 2137–2148 (2018).

39. Zhang, X., Huang, C. T., Chen, J., et al. Pax6 is a human neuroectoderm cell fate determinant. Cell Stem Cell 7, 90–100 (2010).

40. Guo, C., Eckler, M. J., McKenna, W. L., et al. Fezf2 expression identifies a multipotent progenitor for neocortical projection neurons, astrocytes, and oligodendrocytes. Neuron 80, 1167–1174 (2013).

41. Langfelder, P. & Horvath, S. WGCNA: An R package for weighted correlation network analysis. BMC Bioinformatics 9, 1–13 (2008).

42. Milacic, M., Beavers, D., Conley, P., et al. The Reactome Pathway Knowledgebase 2024. Nucleic Acids Res. 52, D672–D678 (2024).

43. Yu, G. & He, Q.-Y. ReactomePA: an R/Bioconductor package for reactome pathway analysis and visualization. Mol. Biosyst. 12, 477–479 (2016).

44. Alvarez, M. J., Shen, Y., Giorgi, F. M., et al. Network-based inference of protein activity helps functionalize the genetic landscape of cancer. Nat. Genet. 48, 838 (2016).

45. Rezsohazy, R., Saurin, A. J., Maurel-Zaffran, C., et al. Cellular and molecular insights into Hox protein action. Development 142, 1212–1227 (2015).

46. Gehring, W. J., Qian, Y. Q., Billeter, M., et al. Homeodomain-DNA recognition. Cell 78, 211–223 (1994).

47. Parker, H. J., Bronner, M. E. & Krumlauf, R. A Hox regulatory network of hindbrain segmentation is conserved to the base of vertebrates. Nature 514, 490–493 (2014).

48. Kirilenko, B. M., Munegowda, C., Osipova, E., et al. Integrating gene annotation with orthology inference at scale. Science 380, eabn3107 (2023).

49. Wang, Q., Van Heerikhuize, J., Aronica, E., et al. Glucocorticoid receptor protein expression in human hippocampus; stability with age. Neurobiol. Aging 34, 1662–1673 (2013).

50. Sukparangsi, W., Morganti, E., Lowndes, M., et al. Evolutionary origin of vertebrate OCT4/POU5 functions in supporting pluripotency. Nat. Commun. 13, 5537 (2022).

51. Zug, R. & Uller, T. Evolution and dysfunction of human cognitive and social traits: A transcriptional regulation perspective. Evol Hum Sci 4, e43 (2022).

52. Berto, S., Mendizabal, I., Usui, N., et al. Accelerated evolution of oligodendrocytes in the human brain. Proc. Natl. Acad. Sci. U. S. A. 116, 24334–24342 (2019).

53. Lowe, C. B. & Haussler, D. 29 mammalian genomes reveal novel exaptations of mobile elements for likely regulatory functions in the human genome. PLoS One 7, e43128 (2012).

54. Kunarso, G., Chia, N.-Y., Jeyakani, J., et al. Transposable elements have rewired the core regulatory network of human embryonic stem cells. Nat. Genet. 42, 631–634 (2010).

55. Wang, J., Xie, G., Singh, M., et al. Primate-specific endogenous retrovirus-driven transcription defines naive-like stem cells. Nature 516, 405–409 (2014).

56. Hsieh, F.-K., Ji, F., Damle, M., et al. HERVH-derived lncRNAs negatively regu-late chromatin targeting and remodeling mediated by CHD7. Life Sci Alliance 5, e202101127 (2022).

57. Ito, J., Sugimoto, R., Nakaoka, H., et al. Systematic identification and characterization of regulatory elements derived from human endogenous retroviruses. PLoS Genet. 13, e1006883 (2017).

58. Osorio, D., Capasso, A., Eckhardt, S. G., et al. Population-level comparisons of gene regulatory networks modeled on high-throughput single-cell transcriptomics data. Nat Comput Sci 4, 237–250 (2024).

59. Enard, W., Khaitovich, P., Klose, J., et al. Intra- and interspecific variation in primate gene expression patterns. Science 296, 340–343 (2002).

60. Oleksiak, M. F., Churchill, G. A. & Crawford, D. L. Variation in gene expression within and among natural populations. Nat. Genet. 32, 261–266 (2002).

61. Romero, I. G., Ruvinsky, I. & Gilad, Y. Comparative studies of gene expression and the evolution of gene regulation. Nat. Rev. Genet. 13, 505–516 (2012).

62. Rohlfs, R. V. & Nielsen, R. Phylogenetic ANOVA: The Expression Variance and Evolution Model for Quantitative Trait Evolution. Syst. Biol. 64, 695–708 (2015).

63. Todd, C. D., Deniz, Ö., Taylor, D., et al. Functional evaluation of transposable elements as enhancers in mouse embryonic and trophoblast stem cells. Elife 8, e44344 (2019).

64. Fuentes, D. R., Swigut, T. & Wysocka, J. Systematic perturbation of retroviral LTRs reveals widespread long-range effects on human gene regulation. Elife 7, e35989 (2018).

65. Halfon, M. S. Perspectives on Gene Regulatory Network Evolution. Trends Genet. 33, 436–447 (2017).

66. Sasaki, K., Yokobayashi, S., Nakamura, T., et al. Robust in vitro induction of human germ cell fate from pluripotent stem cells. Cell Stem Cell 17, 178–194 (2015).

67. Geuder, J., Wange, L. E., Janjic, A., et al. A non-invasive method to generate induced pluripotent stem cells from primate urine. Sci. Rep. 11, 1–13 (2021).

68. Kliesmete, Z., Wange, L. E., Vieth, B., et al. Regulatory and coding sequences of TRNP1 co-evolve with brain size and cortical folding in mammals. Elife 12, e83593 (2023).

69. Kliesmete, Z., Orchard, P., Lee, V. Y. K., et al. Evidence for compensatory evolution within pleiotropic regulatory elements. Genome Res. 34, 1528–1539 (2024).

70. Edenhofer, F. C., Térmeg, A., Ohnuki, M., et al. Generation and characterization of inducible KRAB-dCas9 iPSCs from primates for cross-species CRISPRi. iScience 27, 110090 (2024).

71. Chambers, S. M., Fasano, C. A., Papapetrou, E. P., et al. Highly efficient neural conversion of human ES and iPS cells by dual inhibition of SMAD signaling. Nat. Biotechnol. 27, 275–280 (2009).

72. Ohnuki, M., Tanabe, K., Sutou, K., et al. Dynamic regulation of human endogenous retroviruses mediates factor-induced reprogramming and differentiation potential. Proc. Natl. Acad. Sci. U. S. A. 111, 12426–12431 (2014).

73. Bagnoli, J. W., Ziegenhain, C., Janjic, A., et al. Sensitive and powerful single-cell RNA sequencing using mcSCRB-seq. Nat. Commun. 9, 2937 (2018).

74. Parekh, S., Ziegenhain, C., Vieth, B., et al. zUMIs - A fast and flexible pipeline to process RNA sequencing data with UMIs. Gigascience 7, giy059 (2018).

75. Lun, A. T. L., Bach, K. & Marioni, J. C. Pooling across cells to normalize single-cell RNA sequencing data with many zero counts. Genome Biol. 17, 1–14 (2016).

76. Aran, D., Looney, A. P., Liu, L., et al. Reference-based analysis of lung single-cell sequencing reveals a transitional profibrotic macrophage. Nat. Immunol. 20, 163–172 (2019).

77. Rhodes, K., Barr, K. A., Popp, J. M., et al. Human embryoid bodies as a novel system for genomic studies of functionally diverse cell types. Elife 11, e71361 (2022).

78. Haghverdi, L., Lun, A. T. L., Morgan, M. D., et al. Batch effects in single-cell RNA-sequencing data are corrected by matching mutual nearest neighbors. Nat. Biotechnol. 36, 421–427 (2018).

79. Cannoodt, R., Saelens, W., Sichien, D., et al. SCORPIUS improves trajectory inference and identifies novel modules in dendritic cell development. bioRxiv, 10.1101/079509 (2016).

80. Ahlmann-Eltze, C. & Huber, W. Comparison of transformations for single-cell RNA-seq data. Nat. Methods 20, 665–672 (2023).

81. Van de Sande, B., Flerin, C., Davie, K., et al. A scalable SCENIC workflow for single-cell gene regulatory network analysis. Nat. Protoc. 15, 2247–2276 (2020).

82. Grafen, A. The phylogenetic regression. Philos. Trans. R. Soc. Lond. B Biol. Sci. 326, 119–157 (1989).

83. Housworth, E. A., Martins, E. P. & Lynch, M. The phylogenetic mixed model. Am. Nat. 163, 84–96 (2004).

84. Bininda-Emonds, O. R. P., Cardillo, M., Jones, K. E., et al. The delayed rise of present-day mammals. Nature 446, 507–512 (2007).

85. Christopoulos, D. Introducing unit invariant knee (UIK) as an objective choice for elbow point in multivariate data analysis techniques. SSRN Electron. J. (2016).

86. Corces, M. R., Trevino, A. E., Hamilton, E. G., et al. An improved ATAC-seq protocol reduces background and enables interrogation of frozen tissues. Nat. Methods 14, 959–962 (2017).

87. Vasimuddin, Misra, S., Li, H., et al. Efficient architecture-aware acceleration of BWA-MEM for multicore systems in 2019 IEEE International Parallel and Distributed Processing Symposium (IPDPS) (IEEE, 2019), 314–324.

88. Gaspar, J. M. Genrich: Detecting sites of genomic enrichment version 0.6.1. 2021.

89. Perez, G., Barber, G. P., Benet-Pages, A., et al. The UCSC Genome Browser database: 2025 update. Nucleic Acids Res. 53, D1243–D1249 (2025).

90. Castro-Mondragon, J. A., Riudavets-Puig, R., Rauluseviciute, I., et al. JASPAR 2022: the 9th release of the open-access database of transcription factor binding profiles. Nucleic Acids Res. 50, D165–D173 (2022).

91. Frith, M. C., Li, M. C. & Weng, Z. Cluster-Buster: Finding dense clusters of motifs in DNA sequences. Nucleic Acids Res. 31, 3666–3668 (2003).

92. Paradis, E. & Schliep, K. ape 5.0: an environment for modern phylogenetics and evolutionary analyses in R. Bioinformatics 35, 526–528 (2019).

93. Hoffman, G. E. & Roussos, P. Dream: powerful differential expression analysis for repeated measures designs. Bioinformatics 37, 192–201 (2021).

94. Dobin, A., Davis, C. A., Schlesinger, F., et al. STAR: ultrafast universal RNA-seq aligner. Bioinformatics 29, 15–21 (2013).

95. HUGO Gene Nomenclature Committee (HGNC), European Molecular Biology Laboratory, European Bioinformatics Institute (EMBL-EBI), Wellcome Genome Campus, Hinxton, Cambridge CB10 1SD, United Kingdom. HGNC Database www.genenames.org. Accessed: 2023-5-12.

96. Durinck, S., Moreau, Y., Kasprzyk, A., et al. BioMart and Bioconductor: a powerful link between biological databases and microarray data analysis. Bioinformatics 21, 3439–3440 (2005).

97. McCarthy, D. J., Campbell, K. R., Lun, A. T. L., et al. Scater: pre-processing, quality control, normalization and visualization of single-cell RNA-seq data in R. Bioinformatics 33, 1179–1186 (2017).

98. Cśardi, G. & Nepusz, T. The igraph software package for complex network research. InterJournal Complex Systems 1695, 1–9 (2006).

99. Li, H. Minimap2: pairwise alignment for nucleotide sequences. Bioinformatics 34, 3094–3100 (2018).

100. Langmead, B., Trapnell, C., Pop, M., et al. Ultrafast and memory-efficient alignment of short DNA sequences to the human genome. Genome Biol. 10, R25 (2009).

101. Amemiya, H. M., Kundaje, A. & Boyle, A. P. The ENCODE Blacklist: Identification of Problematic Regions of the Genome. Sci. Rep. 9, 9354 (2019).

102. Huang, X. & Huang, Y. Cellsnp-lite: an efficient tool for genotyping single cells. Bioinformatics 37, 4569–4571 (2021).

103. Huang, Y., McCarthy, D. J. & Stegle, O. Vireo: Bayesian demultiplexing of pooled single-cell RNA-seq data without genotype reference. Genome Biol. 20, 273 (2019).

104. Ritchie, M. E., Phipson, B., Wu, D., et al. limma powers differential expression analyses for RNA-sequencing and microarray studies. Nucleic Acids Res. 43, e47 (2015).

105. Malta, T. M., Sokolov, A., Gentles, A. J., et al. Machine Learning Identifies Stemness Features Associated with Oncogenic Dedifferentiation. Cell 173, 338–354 (2018).

106. Salomonis, N., Dexheimer, P. J., Omberg, L., et al. Integrated Genomic Analysis of Diverse Induced Pluripotent Stem Cells from the Progenitor Cell Biology Consortium. Stem Cell Reports 7, 110–125 (2016).

107. Daily, K., Ho Sui, S. J., Schriml, L. M., et al. Molecular, phenotypic, and sample-associated data to describe pluripotent stem cell lines and derivatives. Sci. Data 4, 170030 (2017).

108. Hao, Y., Stuart, T., Kowalski, M. H., et al. Dictionary learning for integrative, multimodal and scalable single-cell analysis. Nat. Biotechnol. 42, 293–304 (2024).

109. Korsunsky, I., Millard, N., Fan, J., et al. Fast, sensitive and accurate integration of single-cell data with Harmony. Nat. Methods 16, 1289–1296 (2019).

